# Decoding Neoantigen-encoding Tumor-Specific Transcripts Unveils a Shared Target Reservoir for Immunotherapy in Hepatocellular Carcinoma

**DOI:** 10.64898/2025.12.07.692877

**Authors:** Peng Lin, Yifan Wen, Jingjing Zhao, Feifei Zhang, Yaoming Su, Hongwu Yu, Qiaojuan Li, Chengye Liu, Zhixiang Hu, Yan Li, Zhuting Fang, Linhui Liang, Shenglin Huang

## Abstract

**Background and Aims:** Primary liver cancer, predominantly hepatocellular carcinoma (HCC), has limited therapeutic options. While mutation-derived neoantigen vaccine holds promise, its success is hindered by low antigen availability. This study explores transcriptome-derived neoantigens (neoantigen-encoding tumor-specific transcripts, neoTSTs) in HCC, characterizing their features, generation mechanisms, and therapeutic potential.

**Approach and Results:** We analyzed RNA-seq data from 1,013 liver cancer patients and constructed a multi-layered reference dataset. Using a customized pipeline, we identified an average of 60 neoTSTs per patient, significantly surpassing mutation-derived neoantigens (neoMuts). NeoTSTs exhibited higher population frequencies, with 73.1% providing multiple epitopes, and were validated through mass spectrometry and HLA transgenic mouse models. Mechanistically, neoTSTs were generated via retained introns, transposable element activation, HNF4A-regulated alternative promoters, and de novo transmembrane domain (TMD) generation. Single-cell analysis revealed neoTSTs cover >75% of tumor cells and identified antigen-presenting cancer-associated fibroblasts (apCAFs) that enriched in immunotherapy responders and amplified CD4⁺ T-cell responses via MHC-II presentation. In murine HCC models, neoTST vaccination outperformed neoMuts, inducing dual MHC-I/II activation and significant tumor growth inhibition.

**Conclusions:** NeoTSTs represent a superior neoantigen source in HCC, compensating for the limitations of mutation-derived targets. Their abundance, sharedness, and dual MHC pathway activation highlight their potential for personalized immunotherapy, particularly in low-TMB tumors.

**Graphic Abstract:** 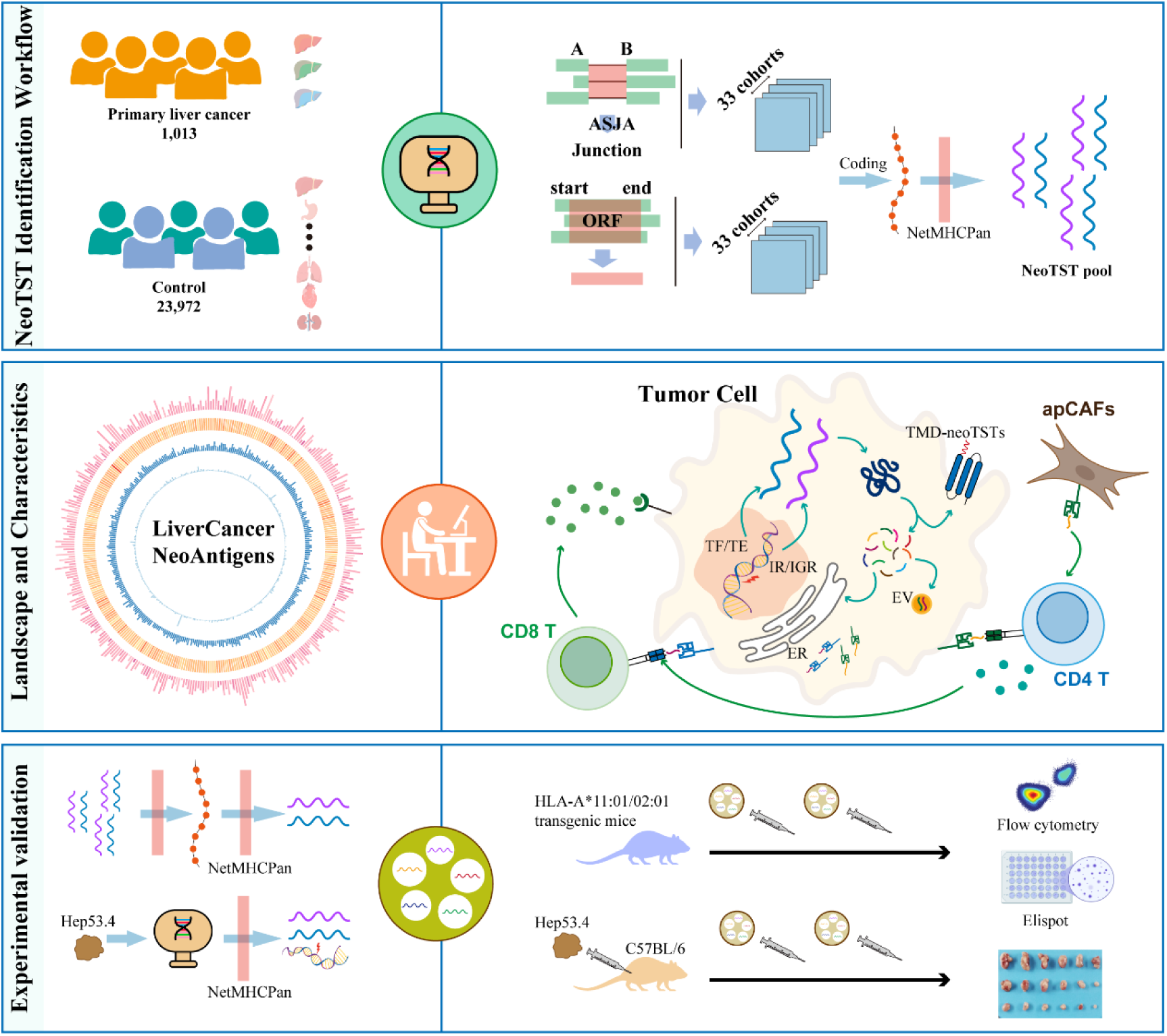

## INTRODUCTION

Primary liver cancer constitutes the third leading cause of cancer-related mortality globally. Hepatocellular carcinoma (HCC), representing approximately 90% of hepatic malignancies, is predominantly diagnosed at advanced stages, conferring poor prognosis (1, 2). Current standard therapies-including surgical resection, liver transplantation, local ablation, and chemotherapy-exhibit significant limitations: chemotherapy demonstrates restricted efficacy with substantial toxicity, while postoperative recurrence occurs in ∼70% of resected patients and 20% of transplant recipients within 5 years (3). For advanced disease, first-line systemic agents (sorafenib, lenvatinib) provide limited clinical benefit (4–6). Notably, recent years have witnessed remarkable advancements in immunotherapy for HCC management (7).

Current advancements in cancer immunotherapy have primarily focused on immune checkpoint inhibitors (ICIs) (8), with multiple clinical trials demonstrating their efficacy as first-line treatments for advanced HCC (9, 10). However, broader application of immune checkpoint inhibitors (ICIs) faces significant challenges in HCC, particularly among patients with immunosuppressive tumor phenotypes (11–13). Most patients derive limited clinical benefit due to insufficient tumor-infiltrating lymphocytes (TILs). Beyond ICIs, tumor-specific neoantigens have emerged as promising targets for cancer immunotherapy (14–17). These uniquely tumor-expressed antigens elicit authentic tumor-specific T-cell responses (18, 19) , induce durable immunity, and minimize therapy-related autoimmunity risks (15). Critically, neoantigen vaccines simultaneously engage CD4⁺ helper T cells and CD8⁺ cytotoxic T cells, reprogramming the tumor microenvironment to convert immunologically “cold” tumors into “hot” ones-thereby overcoming primary resistance to ICIs (20). Additionally, clinical studies have demonstrated the potential of neoantigen vaccines in melanoma, non-small cell lung cancer (NSCLC), and renal cell carcinoma (RCC) (21–23). Notably, recent research has shown that mRNA-based neoantigen vaccines derived from somatic mutations can induce long-term T-cell activation in pancreatic ductal adenocarcinoma (PDAC) (24, 25). Recent advances in HCC neoantigen vaccine research were demonstrated by a Johns Hopkins University School of Medicine clinical trial showing that personalized tumor neoantigen vaccines can potentiate PD-1 inhibitor responses in HCC patients (23). However, most neoantigen vaccines in these studies are mutation-derived, and such neoantigens typically arise from low-frequency somatic mutations, which are rarely shared among patients due to tumor heterogeneity. Moreover, tumors with low mutational burden (e.g., HCC and PDAC) often lack sufficient mutation-derived neoantigen targets, potentially limiting the efficacy of such therapies (26, 27). These findings highlight the limitations of relying solely on mutation-derived neoantigens (neoMuts) in HCC and underscore the urgent need for alternative approaches, such as transcriptome-derived neoantigen discovery, to expand the repertoire of actionable targets.

Recent advances in cancer immunology have revealed that transcriptome-derived neoantigen play a pivotal role in eliciting potent antitumor immune responses (28). Aberrant RNA splicing generates highly immunogenic neoantigens with demonstrated antitumor efficacy in preclinical models (29). These splicing-derived epitopes offer enhanced stability and predictability advantages (30). Our ASJA algorithm (31, 32) has identified clinically relevant tumor-specific transcripts (TSTs) like LIN28B-TST in HCC and MARCO-TST in TNBC (33, 34). Pan-cancer analyses have confirmed TSTs as a promising neoantigen source (27). However, the high complexity of the transcriptome poses significant challenges for validating the specificity of neoantigens. Despite advances in reference datasets, HCC’s transcriptome-originated neoantigens landscape remains underexplored.

To comprehensively identify transcript-derived neoantigens in liver cancer, we conducted a systematic analysis of RNA-seq data from 1,013 liver cancer patients, including 824 HCC, 65 intrahepatic cholangiocarcinomas (ICC), and 124 hepatoblastomas (HB). We established a multi-layered reference dataset comprising 22,734 normal tissue samples, 459 adjacent non-tumor tissues, and 779 non-cancerous liver disease samples. Using a customized RNA-seq-based neoantigen identification pipeline with stringent thresholds for multi-alignment analysis, we systematically identified TSTs (including single- and multi-exonic transcripts) and their corresponding neoantigens. Our analysis revealed an average of 60 neoantigen-encoding TSTs (neoTSTs) per patient, demonstrating significantly greater abundance and coverage compared to neoMuts. Using HLA-A*02:01 and HLA-A*11:01 transgenic mouse models, we validated neoTST-specific CD8^+^ T cell activation. We found that neoTSTs arose from retained introns, transposable element (TE) activation, HNF4A-regulated promoters, and de novo transmembrane domain (TMD) generation. Furthermore, we discovered that antigen-presenting cancer-associated fibroblasts (apCAFs) can activate CD4^+^ helper T cells through neoTST presentation, suggesting a potential therapeutic target for combination immunotherapy. Therapeutic evaluation in murine HCC model demonstrated significant tumor growth inhibition following neoTST vaccination, highlighting its clinical potential.

## METHODS

### Integration of multi-center liver cancer RNA-seq data

To comprehensively characterize the transcriptomic landscape of liver cancer, we integrated RNA-seq data from 1,013 liver tissue samples, including hepatocellular carcinoma (HCC, n = 824), hepatoblastoma (HB, n = 124), and intrahepatic cholangiocarcinoma (ICC, n = 65). The data were aggregated from multiple public repositories and in-house sequencing efforts to ensure broad representation of molecular subtypes and clinical contexts. For HCC, 373 samples were obtained as BAM files from The Cancer Genome Atlas Program (TCGA, https://www.cancer.gov/ccg/research/genome-sequencing/tcga), 105 samples were sequenced internally (raw FASTQ files), and the remaining samples were compiled from 9 independent Gene Expression Omnibus (GEO, https://www.ncbi.nlm.nih.gov/geo) datasets. All 124 HB samples were curated from 7 GEO dataset, while ICC samples included 65 samples from 4 GEO datasets.

For non-TCGA samples, raw FASTQ files were processed using STAR (v2.5.3a) (35) for alignment to the GRCh38/hg38 p12 reference genome in two-pass mode with chimeric junction detection enabled, followed by duplicate marking with Picard Tools. TCGA BAM files were converted to FASTQ using samtools bam2fq and reprocessed identically to ensure uniformity. Transcript abundance was quantified via StringTie (v2.2.1) (36) with GENCODE v29 annotations. This standardized pipeline enabled robust cross-cohort comparisons of liver cancer subtypes.

### Reference transcriptome dataset construction

To establish a comprehensive reference transcriptome dataset for comparative analysis, we integrated RNA sequencing (RNA-seq) data from over 20,000 samples, encompassing normal tissues, adjacent non-tumor tissues, and disease-associated liver tissues across 30 distinct tissue types. The dataset included 1,013 liver tissue samples, 459 adjacent non-tumor samples (of which 50 were paired with tumor samples from our internal cohort), and 22,734 samples from other normal tissues, primarily sourced from the Genotype-Tissue Expression (GTEx v8) project, The Cancer Genome Atlas (TCGA), and Gene Expression Omnibus (GEO) databases. To account for potential confounding effects of non-neoplastic liver diseases, we supplemented the reference with 779 RNA-seq profiles from non-cancer disease tissues (e.g., cirrhosis, hepatitis B/C), curated from 11 independent GEO datasets, 1 ArrayExpress (https://www.ebi.ac.uk/arrayexpress/) dataset: E-MTAB-6863 and 1 in-house data. This diverse dataset ensures robust background signals for tumor-specific analyses while controlling for inflammation- and fibrosis-related transcriptional changes. All data were uniformly processed using the pipeline described in the Integration of multi-center liver cancer RNA-seq data section.

All tissue RNA-seq data were uniformly processed using StringTie for transcript assembly and quantification. Alternative splicing events were identified with ASJA (31), and expression levels were quantified as coverage per million reads (CPT). Subsequently, we constructed a comprehensive reference tissue database by generating 33 independent reference cohorts for each exon junction and single-exon transcript, derived from 30 normal tissue types, adjacent non-tumor tissues, and three liver disease sources. For each single-exon transcript, unique ID was confirmed based on their open reading frame (ORF). For each cohort, three key parameters were calculated: (a) median expression (CPT/TPM), (b) detection frequency, and (c) maximum expression value. This multi-dimensional reference framework enables robust normalization and context-specific analysis of splicing-derived neoantigens.

### TST-derived neoantigens prediction module

To predict TST-derived neoantigens, single-exon and multi-exon TSTs were first merged and filtered based on coding potential, as determined by CPAT (v0.1) (37). Only those transcripts classified as coding by CPAT and containing a complete open reading frame (ORF) were retained as coding TSTs. To minimize false positives, only ORFs of coding TSTs directly resulting from tumor-specific transcriptional events (tumor-specific splicing) and single-exon TSTs were translated into protein sequences (in silico translation) by Biopython (v1.81).

20,420 protein sequences were download from Uniprot as reference protein library. By comparing with the reference protein, the newly identified peptide segments encoded by each transcript were confirmed. For each new peptide segment, 11 amino acids were added at both its beginning and end, resulting in the candidate peptide segment used for predicting potential antigenic epitopes. If the new peptide is at the N-terminus or C-terminus of the protein, then only extend one side of it. HLA class I genotyping was performed using arcasHLA (v3.9) (38) for samples from GEO and in-house data. HLA typing results for TCGA-LIHC samples were obtained from the GDC (see Prediction HLA Typing section). Only HLA alleles supported by NetMHCpan (v4.1) (39) were retained. In total, high-confidence HLA genotypes were determined for 1,048 samples.

NetMHCpan was used to predict 8-12 amino acid peptides binding to HLA, using the parameters ‘-f -inptype 0 -BA -xls -a’. Peptides with predicted binding affinity scores (IC50) <500 nM were classified as either strong binders (SB) or weak binders (WB) and considered candidate neoantigens. Final TST-derived neoantigens were defined as peptides absent in reference protein library.

### Animal model and experimental design

Six- to eight-week-old male C57BL/6 mice (Shanghai Jihui Laboratory Animal Care Co., Ltd.) and genetically modified strains (B6-hHLAA11.1/hB2M and B6-hHLAA2.1/hB2M; GemPharmatech) were used in this study. To establish Hep-53.4 subcutaneous tumors, 2.5×10⁵ cells suspended in 200 μL serum-free DMEM were injected into the right dorsal flank. On day 3 post-inoculation, mice were randomly allocated to three treatment groups: (1) Control: 100 μL PBS administered intramuscularly (i.m.) every 7 days (2 doses total); (2) NeoTSTs: 5 μg neoTST mRNA-LNP (i.m., 100 μL) every 7 days (2 doses); (3) NeoMuts: 5 μg neoMut mRNA-LNP (i.m., 100 μL) every 7 days (2 doses). Tumor dimensions were measured every 2-3 days using digital calipers, with volumes calculated as V = (L × W²)/2. A strict endpoint criterion (tumor volume >1,500 mm³ or 20 mm in any dimension) was enforced. Mice were housed under specific pathogen-free conditions (12-h light/dark cycle, 20-22°C, 40-60% humidity). All procedures complied with Shanghai Laboratory Animal Care Association guidelines and were approved by the Fudan University Institutional Animal Care and Use Committee (IACUC protocol: 202510FD0002).

### Statistical analyses

Statistical analyses were performed using R (v4.0.2) and python (python 3.9.18). Comparisons between two groups were assessed by the Wilcoxon rank-sum test. Survival analyses utilized Kaplan-Meier estimators with log-rank tests to determine statistical significance. One-way ANOVA was employed for group comparisons in tumor size and flow cytometry data; results are presented as mean ± SEM. Significance levels were defined as follows: *P < 0.05, **P < 0.01, ***P < 0.001.

Full and detailed descriptions of research materials and experiment methods are provided in online Supporting Information.

## RESULTS

### Landscape of neoantigen-encoding tumor-specific transcripts (neoTSTs) in liver cancer

RNA-seq data from 1,013 liver cancer samples were analyzed to comprehensively characterize transcript-derived neoantigens in liver cancer (Figure 1A, Supplemental Figure S1A). To rigorously identify tumor-specific transcripts (TSTs), we constructed a comprehensive reference dataset consisting of 33 independent reference cohorts, which included over 20,000 normal tissue samples (29 tissues), 459 adjacent non-tumor tissues, and 779 non-cancer liver disease samples, encompassing nearly 3 million splice junctions (Figure 1A, Supplemental Figure S1A-B). We developed a robust computational pipeline for neoantigen discovery (Figure 1A, Methods): (1) Transcript assembly: RNA-seq data from tumor and control samples were aligned using STAR and assembled into transcripts with StringTie. (2) Reference construction: Comprehensive reference datasets were built for both multi-exonic and single-exonic transcripts. (3) TST detection: Tumor samples were compared against controls using stringent thresholds to identify TSTs under multi-exonic and single-exonic conditions. (4) Neoantigen prediction and evaluation: Open reading frames (ORFs) were predicted computationally to assess coding potential and identify novel peptides. Potential epitopes were predicted using NetMHCpan (v4.1) with affinity evaluation. This integrated pipeline enables systematic identification of neoTSTs and their potential epitopes in liver cancer.

**Figure 1.**
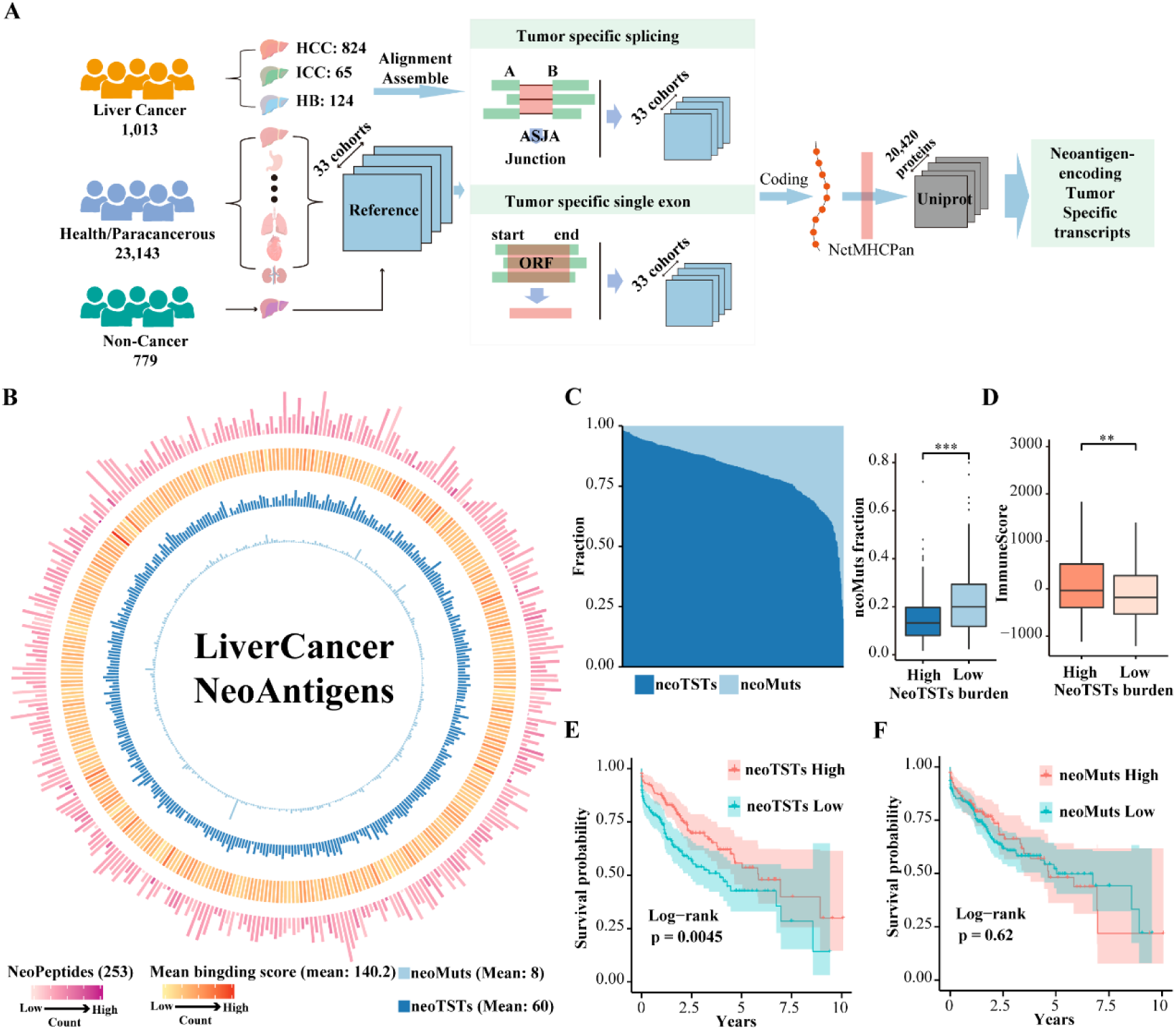
Landscape of Neoantigen-encoding tumor-specific transcripts (neoTSTs) in 1,013 Liver Cancer. (A) Workflow for identifying neoTSTs: (1) multicenter data collection and construction of reference datasets; (2) detected tumor-specific transcripts (TSTs) separately for single-exon and multi-exon transcripts; (3) neoantigen prediction from coding TSTs. (B) Circos plot summarizing neoantigen features per liver cancer sample: (1) number of TST-derived neoantigens; (2) binding affinity scores of TST-derived neoantigens; (3) number of neoantigen-encoding TSTs (neoTSTs); and (4) number of mutation-derived neoantigens (neoMuts). (C) Distribution and comparative analysis of neoTST and neoMut burden in patients. Left panel: Stacked bar plot displaying the relative proportion of neoTSTs (dark blue) and neoMuts (light blue) for each individual patient. Right panel: Boxplot comparing neoMut proportions between high vs. low neoTST-burden group. Statistical significance was determined by Wilcoxon rank-sum test (***p < 0.001). (D) The boxplot demonstrates significantly higher immune scores (calculated via ESTIMATE algorithm) in patients with high neoTST burden compared to those with low neoTST burden. Statistical significance was determined by Wilcoxon rank-sum test (**p < 0.01). (E) Kaplan-Meier survival analysis stratified by neoTST burden (high vs. low) in TCGA-LIHC cohort. Log-rank test was used.The optimal cutoff point of TST-derived neoantigen burden was applied. The mean HLA-C associated neoTST burden was used for stratification. (F). Kaplan-Meier survival curves for the TCGA-LIHC cohort stratified by neoMuts burden. The mean neoMut burden was used for stratification.

Approximately 150,000 splice junctions were detected per patient by ASJA (Supplemental Figure S1C). Through stringent filtering criteria, we identified an average of 300 TSTs per tumor sample, which were further refined to ∼60 neoTSTs per case (44 multi-exonic transcripts (METs) and 16 single-exonic transcripts (SETs); Figure 1B, Supplemental Figure S1D)). In stark contrast, only 8 neoMuts were identified per patient on average, highlighting the quantitative superiority of neoTSTs. Immunogenicity prediction revealed that each neoTST generated ∼5 immunogenic peptides (253 NeoPeptides/patient on average) with a mean HLA-I affinity score of 140.2, demonstrating high binding potential. Notably, neoTST and neoMut burdens exhibited a complementary pattern: patients with low neoMuts loads showed significantly elevated neoTST levels (Figure 1C), suggesting that neoTSTs effectively compensate for the paucity of immunogenic epitopes in TMB-low tumors.

Comparative analysis revealed that the number of candidated neoTSTs identified was remarkably consistent across the 3 cancer types (Supplemental Figure S1D). Despite substantial inter-patient and inter-tumor heterogeneity, neoTSTs exhibited significant overlap between cancer types, with 24.6% of HB-associated neoTSTs and 27.5% of ICC-associated neoTSTs being shared with HCC (Supplemental Figure S1E). Given HCC’s predominance among liver cancers, subsequent analyses focus specifically on this malignancy. Our analysis of 824 HCC cases identified 347 patients (42.1%) with detectable HBV infection (Supplemental Figure S1F). Comparative analysis demonstrated that HBV-positive patients exhibited a significantly enriched repertoire of transcript-derived neoantigens (p < 0.05) (Supplemental Figure S1G). These findings suggest that HBV infection may actively promote neoantigen generation through the formation of neoTSTs, potentially via virus-induced genomic instability or transcriptional dysregulation.

We evaluated the immune activation score for each patient and found that those with a high neoTST burden exhibited significantly higher immune scores, suggesting stronger immune activation (Figure 1D). Given that immune activation levels are often associated with improved clinical outcomes, we further analyzed the survival correlation of neoPeptide burden across different HLA subtypes, accounting for population heterogeneity and HLA preferences. While the overall neoTST burden showed no clear prognostic association, distinct patterns emerged in specific populations: in European/American cohorts, patients with high HLA-C neoPeptide loads demonstrated significantly better survival outcomes, whereas in Asian cohorts, HLA-A emerged as the dominant prognostic marker (Figure 1E, Supplemental Figure S1H-I). In contrast, neoMuts showed no significant correlation with patient prognosis (Figure 1F). These findings highlight population-specific immunological heterogeneity and suggest that HLA-restricted neoTST presentation may serve as a more robust biomarker for immune responsiveness than conventional mutation-based neoantigens.

### Mass spectrometry and immunopeptidomics-based identification and immunogenicity assessment of neoTSTs in liver cancer

Compared to neoMuts (mean frequency < 0.001), neoTSTs exhibited significantly higher population frequencies (mean frequency > 0.02), suggesting their potential as shared neoantigens capable of broader patient coverage (Figure 2A, Supplemental Figure S2A). Independent analysis of the top 20 neoTST candidates demonstrated their broad patient applicability, directly targeting 708 cases. This demonstrates both the polyvalent nature of neoTSTs and the existence of shared epitopes across different neoTSTs, making the identification of identical neoTSTs in unmatched samples a feasible prospect. To validate the translational potential and population-wide sharedness of neoTSTs, we analyzed mass spectrometry data from 159 paired HCC and adjacent non-tumor samples (Figure 2B). Peptide-level evidence was detected for 1,558 (3.9%) neoTSTs across the cohort, alongside 12,911 UniProt-annotated proteins (Figure 2C, Supplemental Figure S2B). Notably, specific neoTSTs such as LIN28B-neoTST and TSPAN13-neoTST were directly verified at the protein level (Figure 2D-F). Mass spectrometry analysis confirmed the high tumor specificity of neoTSTs (Figure 2G). 73.1% of neoTSTs provided ≥2 predicted epitopes matching patient-specific HLA haplotypes, with 59.4% capable of binding multiple HLA allelic subtypes (self-HLA), demonstrating superior immunogenic potential compared to neoMuts that typically yield single epitopes (Figure S2A). Further analysis of immunopeptidomic profiles from 15 HCC patients revealed ≈ 3 HLA-I immunogenic neoTSTs per patient after stringent filtering, with two-thirds originating from novel splice junctions (Supplemental Figure S2C). Representative examples like G5TA1-neoTST (Figure 2H, I) confirmed both the translational capacity and immunogenic superiority of neoTSTs. Analysis of HLA restriction patterns for immunopeptidome-validated neoantigens revealed significant allelic bias, with HLA-A*02:01 and HLA-B*07:02 emerging as dominant restriction element (Supplemental Figure S2D).

**Figure 2.**
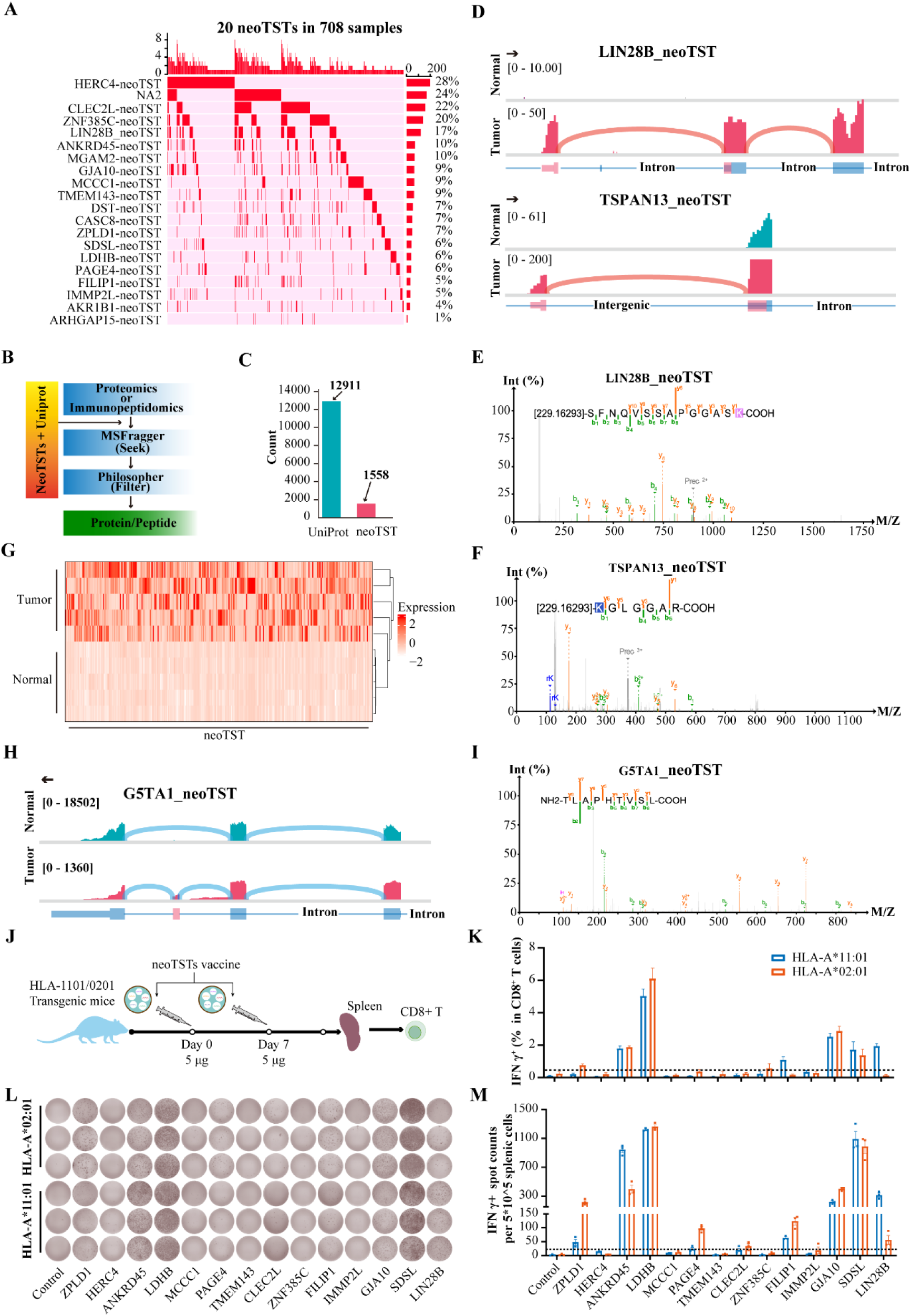
Mass Spectrometry and Immunopeptidomics-Based Identification and Immunogenicity Assessment of neoTSTs in Liver Cancer. (A) Distribution and expression frequency of 20 neoTSTs across samples. The waterfall plot illustrates the presence (red) or absence (white) of 20 neoTSTs (left panel) in the analyzed cohort. Each column represents an individual sample, and each row corresponds to a specific neoTST. Red indicates detectable expression of the neoantigen. (B) Overview of the proteomic and immunopeptidomic analysis pipeline. (C) The bar plot compares the total number of UniProt background proteins and neoTST-derived proteins detected across samples. Values represent the cumulative protein counts from all replicates/experimental conditions. (D) Sashimi plot of LIN28B-neoTST and TSPAN13-neoTST. (E-F) Mass spectrometry (MS) spectrum of peptides of LIN28B-neoTST and TSPAN13-neoTST. (G) The heatmap displays normalized expression levels of neoTST-derived peptides identified by mass spectrometry, comparing tumor and matched normal adjacent tissues. Each column represents the expression of each neoTST in a single experiment. A single experiment involves 5 tumor samples and 5 corresponding adjacent tissue samples. (H) Mass spectrometry (MS) spectrum of peptide of G5TA1-neoTST. (I) Sashimi plot of G5TA1-neoTST. (J) On the 10th day after the second vaccination, spleen cells were collected and mixed with individual neoTST mRNA for in vitro culture. The process was monitored using the ELISpot and flow cytometry assay. (K) Flow cytometry quantification of IFN-g-secreting CD8^+^T cells derived from HLA-A*02:01 and HLA-A*11:01 transgenic mice after stimulation with neoTST mRNA-LNP vaccine. (L) ELISpot images of IFN-g-secreting spleen cells derived from HLA-A*02:01 and HLA-A*11:01 transgenic mice after stimulation with neoTST mRNA-LNP vaccine. (M) ELISpot quantification of IFN-g-secreting spleen cells derived from HLA-A*02:01 and HLA-A*11:01 transgenic mice after stimulation with neoTST mRNA-LNP vaccine.

To validate endogenous immunogenicity, we selected 14 neoTSTs (Supplemental Table S1) and conducted further experimental verification in HLA-A*02:01 and HLA-A*11:01 transgenic mice. Each candidate fragment contained at least one HLA-A*11:01- or HLA-A*02:01-restricted epitope, with seven neoTSTs fragments harboring binding epitopes for both HLA subtypes. We designed and produced a mixed neoTST mRNA-LNP vaccine, which was administered via two intramuscular injections (5 μg/dose) at a 7-day interval to humanized transgenic mice expressing either HLA-A*11:01 or HLA-A*02:01. One week after immunization, splenocytes were subjected to IFN-γ ELISpot and flow cytometry to quantify antigen-specific T-cell responses (Figure 2J). The flow cytometry results showed that 50% (7/14) of the neoTSTs elicited robust HLA-A*11:01- or HLA-A*02:01-restricted T-cell activation, including LDHB, GJA10, ANKRD45, SDSL, ZPLD1, FILIP1 and LIN28B-neoTST (Figure 2K). Moreover, 57.1% (4/7) of these activated T-cell responses were specific to both HLA subtypes transgenic mice. Notably, all flow-cytometry-positive neoTSTs were confirmed identified by ELISPOT assay, which additionally included the T-cell response triggered by PAGE4-neoTST (Figure 2L, M). The four double-positive neoTSTs exhibited particularly strong responses, with consistent patterns across mice, demonstrating the reliability of these findings and supporting neoTSTs as promising universal antigen candidates.

### Retention of noncoding sequences and transposon-driven activation generate high-yield neoantigen expansion

Genomic annotation revealed that 96.6% of neoTSTs originate from unannotated splicing sites (Figure 3A), highlighting the critical role of alternative mechanisms in neoantigen generation. To systematically characterize splicing diversity, we classified aberrant splicing events into four categories: exon skip (ES), exon truncation (ET), intron retention (IR), and intergenic region retention (IGR) (Supplemental Figure S3A). Quantitative analysis demonstrated that neoTSTs predominantly arise from ET events (68%), followed by IR (18%), IGR (6%) and ES (6%), with only a minor contribution from IGR (2%) (Supplemental Figure S3B). Representative examples of ET (e.g., DOCK7-neoTST), and IR (e.g., ZPLD1-neoTST) and IGR (e.g., IMMP2L-neoTST) events are illustrated in Figure 3C. Notably, IR and IGR events exhibited superior neoantigen-generating capacity, producing 2.3-fold more immunogenic peptides compared to single-site anomalies. Multi-site aberrant events further enhanced neoantigen yield (Figure 3B), attributable to their truncated open reading frames (ORFs) (Figure 3D). These shortened ORFs confer dual advantages: (1) accelerated ribosomal initiation and elongation (40), and (2) enhanced peptide processing and TAP-dependent transport (41), collectively optimizing neoantigen presentation. While most splicing junctions adhered to canonical GT-AG signals (Supplemental Figure S3C), exon-exon splicing exhibited widespread non-canonical signal selection (e.g., GC-AG, AT-AC; Figure 3E). These findings establish that beyond coding-region frameshifts (ET/ES), non-coding sequence retention (IR/IGR) serves as a potent neoantigen source, with their truncated ORFs potentially enhancing immunogenicity through streamlined biogenesis and presentation pathways.

**Figure 3.**
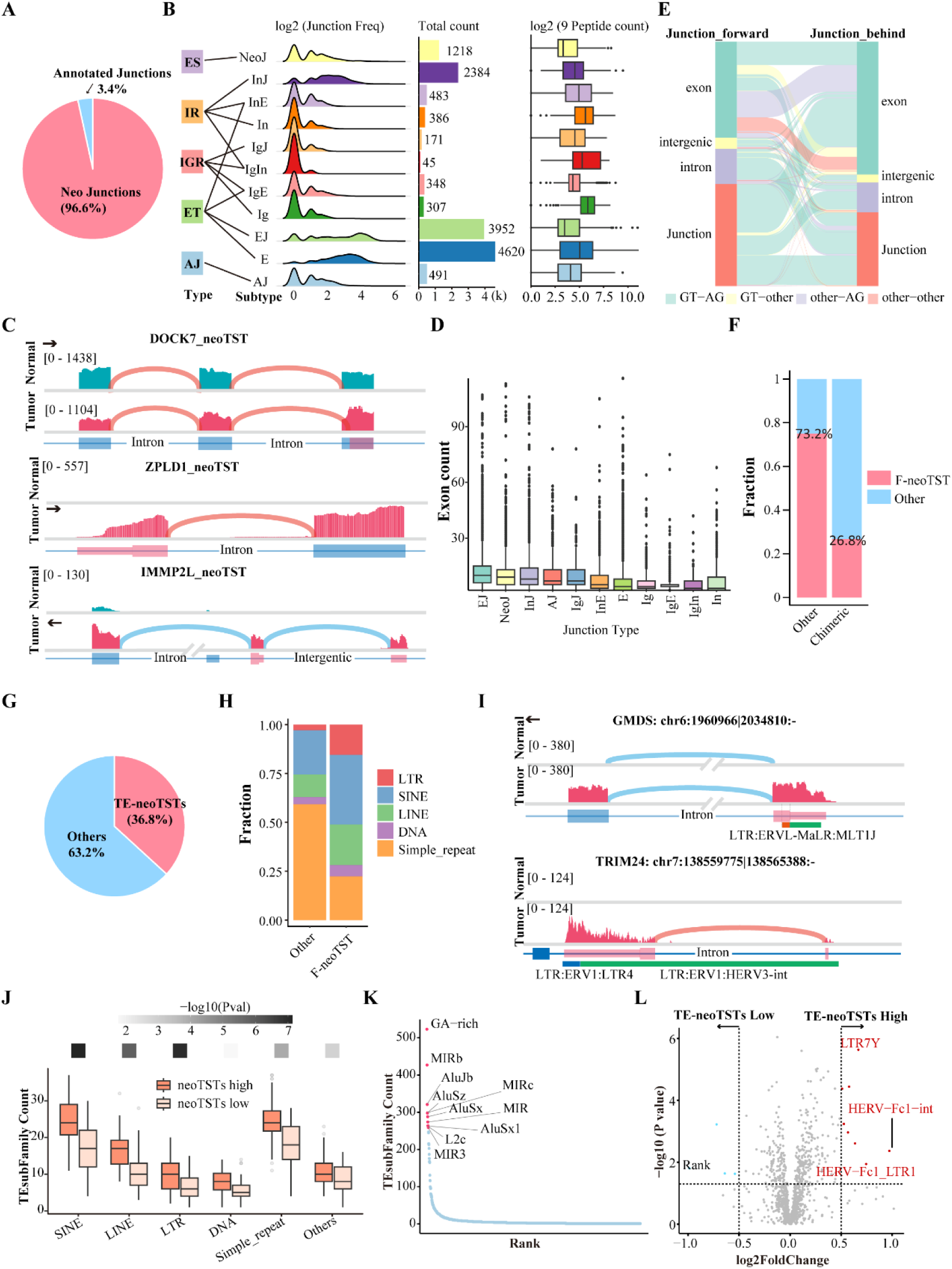
Retention of noncoding sequences and transposon-driven activation generate high-yield neoantigen expansion. (A) The pie chart illustrates the proportion of annotated (colored) versus unannotated (grey) tumor-specific splicing (TSS) identified in liver cancer. (B) Characterization of five major tumor-specific splicing types and their 11 reference genome location sources. Left panel: Density plot showing the distribution frequency. The x-axis represents TSS classification, and the y-axis reflects normalized frequency. Middle panel: Bar plot displaying the total count of TSS events. Right panel: Boxplot comparing the number of 9-mer peptides from neoTSTs for each TSS type, the x-axis reflects count of 9-mer peptides after the log2 transformation. ES: exon skipping, ET: exon truncation, IR: intron retention, IGR: intergenic region retention, AJ: annotated junction. The sources of the 11 genomic locations are specifically described in the method. (C) Sashimi plots of DOCK7 (ET), ZPLD1 (IG), and IMMP2L (IGR). (D) The boxplot compares the number of exons contained within neoTSTs generated by distinct splicing event subtypes. (E) The Sankey diagram illustrates the distribution of splicing signals (colored links) derived from different reference genome locations (left nodes). Non-canonical splicing signals (colored in non-green hues) are contrasted with canonical GT-AG signals (green). Transparent link opacity emphasizes signal flow density, with node widths proportional to the frequency of genomic region contributions. (F) Bar plot compares the percentage of F-neoTSTs (functional neoantigen transcripts) within chimeric neoTSTs (fusion-derived, left bar) and non-chimeric neoTSTs (canonical splicing-derived, right bar). (G) The proportions of TE-neoTSTs and non-TE-neoTSTs. (H) The stacked bar plot compares the distribution of transposable element (TE) subfamilies (e.g., LINE, SINE, LTR, DNA transposons) between F-neoTSTs (functional, left bars) and non-F-neoTSTs (non-functional, right bars) derived from TE-associated splicing events (TE-neoTSTs). Subfamilies are color-coded, and the y-axis represents the percentage of total TE-neoTSTs in each group. (I) Sashimi plot of 2 TE-neoTST: GMDS-neoTST and TRIM24-neoTST. (J) The boxplot compares the expression levels of individual TE classes between high neoTSTs burden samples and low neoTSTs burden samples. Each box represents the distribution of subfamily expression within a TE type. Dots indicate outliers. (K) Number of neoTSTs derived from individual TE subfamilies, ordered by descending count (left to right on the x-axis). The y-axis represents the absolute count of neoTSTs per subfamily. (L) The volcano plot displays log2 fold-change (x-axis) and statistical significance (-log10 adjusted p-value, y-axis) of TE subfamily expression in high versus low TE-neoTST burden groups.

Alternative promoter usage through transcription start site (TSS) selection represents a significant source of neoTST generation (42). Our analysis revealed that 67.9% of neoTSTs consisted entirely of novel exons (non_chimeric-neoTSTs), while 32.1% were chimeric transcripts (chimeric-neoTSTs) (Supplemental Figure S3D-E). Notably, 26.8% of chimeric-neoTSTs utilized novel TSS loci (F-neoTSTs), suggesting their regulation by alternative promoters (Figure 3F). Amino acid position analysis of NeoPeps demonstrated strong enrichment at N-terminal chimeric regions (Supplemental Figure S3F), further supporting alternative promoter activity as a key driver of neoTST biogenesis. Given the established role of transposable elements (TEs) in generating aberrant splicing isoforms (26) and their documented promoter activity in cancer (43, 44), we systematically analyzed TE content within the first exon and 200 bp upstream region of neoTSTs. Strikingly, 36.8% of neoTSTs harbored TE sequences in these regulatory regions (Figure 3G). F-neoTSTs exhibited significant enrichment of LTR (∼6-fold), SINE (∼1.5-fold), and LINE (∼2-fold) elements compared to background, with LTR elements showing particularly strong association (Figure 3H, I). These findings align with prior reports of LTR-specific promoter activity (45, 46) and collectively demonstrate that TEs - especially LTR, SINE, and LINE elements - may function as alternative promoters to activate neoTSTs transcription in cancer.

To substantiate whether TEs drive neoTSTs expression, we quantified TE expression levels across all HCC patients. Analysis revealed significantly elevated TE expression in patients with high neoTSTs burden (*p* < 0.001, Figure 3J). Characterization of TE-associated neoTSTs subtypes identified distinct TE subfamily enrichments: GA-rich elements predominated in non-F-neoTSTs, while Alu and MIR elements were enriched in F-NeoTSTs (Figure 3K, Supplemental Figure S3G). Differential expression analysis further demonstrated that upregulated TEs clustered predominantly in high-neoTST-burden samples (Figure 3L). Notably, LTR7Y expression exhibited a proportional correlation with neoTST burden (*p* = 1.2e-7, Supplemental Figure S3H). These findings collectively demonstrate that TE-specific activation may provide alternative promoters that potentiate neoTSTs expression in HCC. In addition to regulating the transcription of neoTSTs, amino acid sequences derived from TEs can also serve as a substantial source of neo-peptides (neoPeps). We characterized TE-chimeric neoTSTs (Methods) and found that 21.4% of neoTSTs were TE-neoTSTs, generating NeoPeptides derived from TE sequences (Supplemental Figure S3I).

### neoTSTs could generate de novo transmembrane domains (TMDs)

The generation of immunogenic neoantigens requires proteolytic processing and TAP-mediated transport of peptides into the endoplasmic reticulum (ER) for MHC-I loading, a process potentially facilitated by transmembrane domains (TMDs). Our analysis revealed that approximately 15% of neoTSTs contained TMDs, with 67% representing novel transmembrane domains acquired through chimeric transcript formation (Figure 4A-B). While wild-type membrane proteins like ZPLD1-neoTST, HERC4-neoTST and GJA10-neoTST retained their original TMDs after alternative splicing (Figure 4C), non-membrane proteins such as SDSL-neoTST gained de novo transmembrane capability through structural reorganization (Figure 4D). Strikingly, despite significant alternative splicing-induced modifications, all TMD-containing neoTSTs maintained functional transmembrane topology (Figure 4E, Supplemental Figure S4A). Immunofluorescence validation confirmed membrane localization not only for TMD-neoTSTs derived from native membrane proteins (ZPLD1-neoTST, HERC4-neoTST, GJA10-neoTSTs) but also for SDSL-neoTST, demonstrating the widespread acquisition of membrane-targeting capacity among neoTSTs (Figure 4F). In vivo validation showed three of four tested membrane-capable neoTSTs (including the de novo transmembrane SDSL-neoTST and multi-TMD GJA10-neoTST) potently activated HLA-restricted T cells (Figure 2I-K).

**Figure 4.**
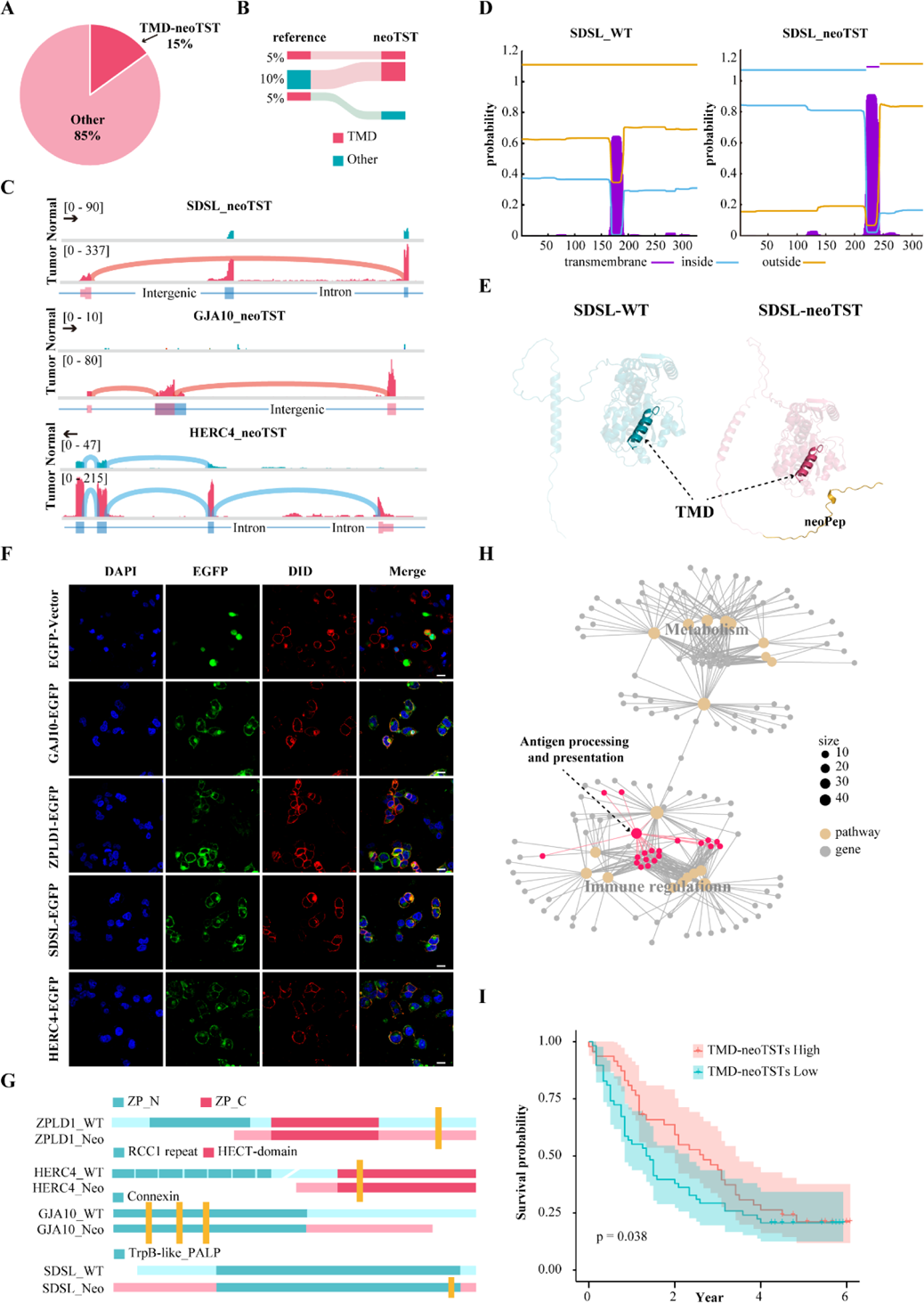
neoTSTs could generate de novo transmembrane domains (TMDs). (A) The proportions of TMD-neoTSTs. (B) Transmembrane domain (TMD) dynamics between reference proteins and neoTST-derived isoforms. The Sankey diagram visualizes changes in TMD status (presence/absence) between reference protein isoforms (left nodes) and neoTST-derived isoforms (right nodes). Node colors: Red = TMD present, Green = TMD absent. Link colors: TMD gained or retained in neoTSTs, Green = TMD lost in neoTSTs. Link widths correspond to the frequency of each transition. (C) Sashimi plot of 3 TMD-neoTST: SDSL-neoTST, GJA10-neoTST and HERC4-neoTST. (D) TMD prediction profiles along the protein sequence (x-axis: residue position; y-axis: probability). Purple: predicted transmembrane domains (probability ≥ 0.7); blue line: cytoplasmic regions; yellow line: extracellular regions (E) The figure presents the 3D structural models of: Wild-type SDSL (blue ribbon diagram) SDSL-neoTST (red ribbon diagram). (F) The panel displays immunofluorescence images of four TMD-neoTSTs (rows), each visualized with four markers (columns): DAPI (blue): nuclear staining. GFP (green): TMD-neoTST fusion protein expression. DID (red): membrane dye (labels plasma membrane or organelles). Merge: overlay of all channels. (G) Domain annotation of 4 TMD-neoTSTs and their wild-type proteins. yellow vertical bars: TMDs. (H) The network diagram illustrates functional associations between wild-type proteins (gray nodes) and their enriched biological pathways (yellow nodes), with antigen processing and presentation pathways highlighted in red. (I) Kaplan-Meier survival curves for the in-house cohort stratified by TMD-neoTST burden. The mean TMD-neoTST burden was used for stratification.

Functional domain annotation revealed that TMD-neoTST-derived membrane domains frequently occupied critical protein regions (Figure 4G), suggesting preserved functional capacity. Notably, most membrane-capable neoTSTs originated from originally non-transmembrane proteins (Figure 4B). Enrichment analysis demonstrated significant association with antigen processing/presentation pathways alongside metabolic functions (Figure 4H). Clinically, patients with higher TMD-neoTST burden exhibited improved survival (Figure 4I), suggesting that acquired transmembrane functionality may enhance neoantigen presentation efficiency and ultimately improve patient outcomes through structural and functional modulation of antigen processing.

### HNF4A binding activates neoTSTs transcription in HCC

To further explore potential regulatory mechanism of neoTSTs, we performed transcription factor enrichment analysis on the first exon regions of F-neoTSTs. Our results revealed significant enrichment of transcription factors such as HNF4A and HNF4G in the initiation regions of F-neoTSTs. Notably, HNF4A was found to bind to the promoter regions of 51.03% of F-neoTSTs, and its binding level showed a significant positive correlation with neoTST burden (Figure 5A). For instance, both PAGE4 and CASC8-neoTSTs contained HNF4A-binding sites within their promoter regions (Figure 5B). Furthermore, analysis of bulk RNA-seq data confirmed that HNF4A expression was positively correlated with neoTST burden, with HNF4A-high patients exhibiting a greater number of neoTSTs (Figure 5C). Single-cell RNA-seq data with 140 HCC samples across 20 datasets (47) were integrated to further elucidate the regulatory mechanism of HNF4A on neoTSTs. We quantified neoTSTs in the tumor cells of each patient and observed that the expression of HNF4A was highly specific to liver cancer cells and showed significant co-localization with neoTSTs expression (Figure 5D-F, Supplemental Figure 5A). Statistical analysis further confirmed a strong positive correlation between HNF4A expression and neoTSTs burden at the single-cell level (Figure 5G), supporting a functional link between them.

**Figure 5.**
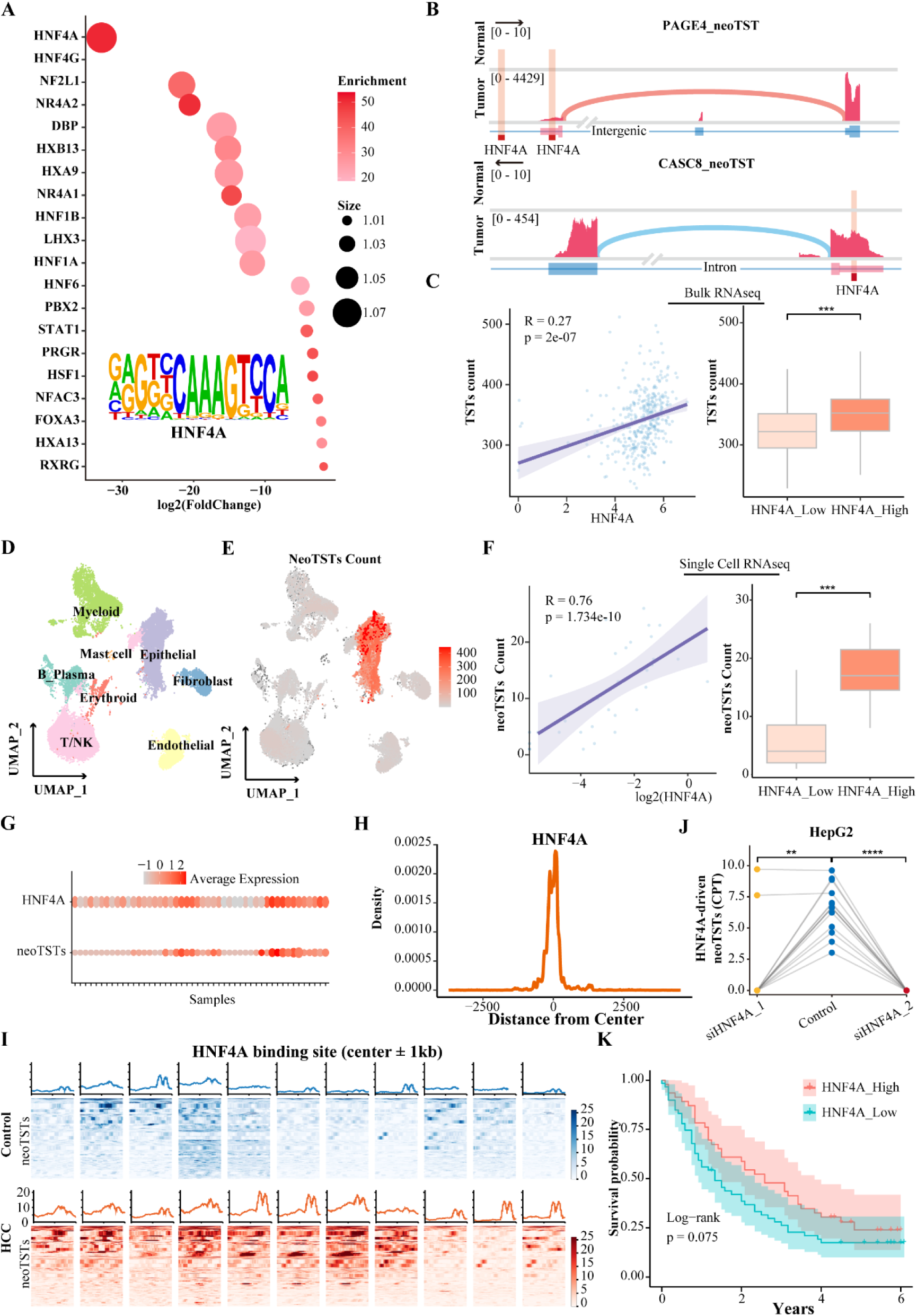
HNF4A binding activates neoTSTs transcription in HCC. (A) The bubble plot illustrates transcription factor (TF) enrichment in the transcriptional start regions of F-neoTSTs, with the most highly enriched TF, HNF4A, emphasized. Bubble color intensity reflects enrichment significance (darker = stronger), and size represents fold enrichment relative to background. Inserted below the plot is the sequence logo of the HNF4A binding motif at F-neoTST promoters, showing conserved nucleotides with letter height indicating sequence conservation. (B) Sashimi plot of 2 HNF4A binding neoTSTs: PAGE4-neoTST and CASC8-neoTST. (C) The panel combines two analyses of bulk RNA-seq data to link HNF4A expression with neoTST load. Left: Scatter plot showing the positive correlation between HNF4A expression (x-axis, TPM) and neoTST burden (y-axis, count per sample). A trendline (blue dashed line) indicates a significant linear relationship (r = 0.27, p < 0.001). Right: Boxplot comparing neoTST burden between samples stratified by HNF4A expression (high vs. low, defined by mean split). The boxplot shows significantly higher neoTST burden in the HNF4A-high group compared to the HNF4A-low group. Statistical significance was determined by Wilcoxon rank-sum test (***p < 0.001). (D) The UMAP (Uniform Manifold Approximation and Projection) plot displays clustering of single cells (dots) from liver cancer tissues, colored by their cell type annotations. Each cluster corresponds to a distinct cell population, with major types labeled (including epithelial, Myeloid, T/NK, Erythroid, B Plasma, Mast cell, Fibroblast). (E) The UMAP plot highlights neoTSTs count within the single-cell landscape of liver cancer. (F) The panel combines two analyses of single cell RNA-seq data to link HNF4A expression with neoTST load. Left: Scatter plot showing the positive correlation between HNF4A expression (x-axis, log2(normalized count)) and neoTST burden (y-axis, count per sample). A trendline (blue dashed line) indicates a significant linear relationship (r = 0.76, p < 0.001). Right: Boxplot comparing neoTST burden between samples stratified by HNF4A expression (high vs. low, defined by mean split). The boxplot shows significantly higher neoTST burden in the HNF4A-high group compared to the HNF4A-low group. Statistical significance was determined by Wilcoxon rank-sum test (***p < 0.001). (G) The dot plot displays the relationship between HNF4A expression and neoTST burden at the single-cell RNA-seq data across patient samples. Rows: Individual patient samples. (H) The schematic illustrates putative HNF4A binding motifs within a 6kb window (x-axis: position relative to the transcriptional start site [TSS] of F-neoTSTs; range: -3kb to +3kb from TSS, marked by vertical dashed line). (I) ATAC-seq signals (±1kb around HNF4A binding sites) between 11 HCC tumors (red) and 11 normal liver tissues (blue), aligned to the center of HNF4A peaks (x-axis: genomic position; y-axis: normalized ATAC-seq signal intensity). (J) HNF4A-driven neoTST expression (CPT) in HepG2 between HNF4A knockdown group (siHNF4A_1, siHNF4A_2) and non-knockdown group (Control). Statistical significance was determined by Wilcoxon rank-sum test (**p < 0.01, ****p < 0.0001). (K) Kaplan-Meier survival curves for the in-house cohort stratified by HNF4A expression. The mean FPKM of HNF4A was used for stratification.

The analysis revealed a significant central enrichment phenomenon at the binding site of HNF4A in the promoter region of F-neoTSTs. (Figure 5H). To investigate whether HNF4A regulates neoTSTs coupling with epigenetic mechanisms, we analyzed ATAC-seq data from 22 internal samples (11 controls and 11 HCC cases). The results revealed that potential HNF4A binding sites exhibited significantly increased chromatin accessibility in HCC patients, with these sites predominantly located in non-coding regions such as promoters and enhancers. This suggests that tumor-specific epigenetic opening may recruit HNF4A binding, thereby activating neoTSTs transcription (Figure 5I). To further validate HNF4A’s regulation of neoTST biogenesis, we analyzed RNA-seq data from HepG2 cells following HNF4A knockdown (GSE199068, two independent siRNA constructs). Both knockdown conditions showed significant reductions in HNF4A-driven neoTSTs (Figure 5J), establishing a causal relationship between HNF4A expression and neoTST generation. Survival analysis further indicated that high HNF4A expression was associated with favorable patient prognosis (Figure 5K).

### Heterogeneity of neoTSTs at single-cell resolution

HCC exhibits extensive tumor heterogeneity and a complex immune microenvironment, necessitating analysis of neoTST heterogeneity at single-cell resolution. We collected single-cell RNA sequencing data from a cohort of 112 liver cancer patients and 15 healthy donors, and developed a computational pipeline for quantifying and screening neoTSTs at the single-cell level (Figure 6A). After reclustering, we annotated patient cells into 16 distinct cell types, including tumor cells (HCC and ICC tumor cells), 12 immune cell subtypes, and two stromal cell types (Figure 6B). We observed that neoTSTs are primarily produced by tumor cells, enriched in tumor epithelial cells (Figure 6C). Although the proportion of cancer cells expressing neoTSTs varied among patients, and the specific neoTSTs expressed differed across cancer cells, neoTSTs collectively covered over 75% of HCC cells in the cohort, with the majority of patients showing neoTST expression in more than 75% of their tumor cells (Figure 6D). These findings suggest that personalized vaccine design should target multiple neoTST antigens to achieve optimal therapeutic efficacy. To further investigate cellular heterogeneity, we stratified patients into low- and high-neoTST burden groups based on the proportion of tumor cells expressing neoTSTs (Figure 6E). Analysis of T cell composition showed that the proportion of CD8^+^ T cells in the high-burden group was more than double that in the low-burden group, indicating a heightened state of immune activation in high-neoTST tumors (Figure 6F). These results further support the role of neoTSTs in promoting immune activation in HCC patients.

**Figure 6.**
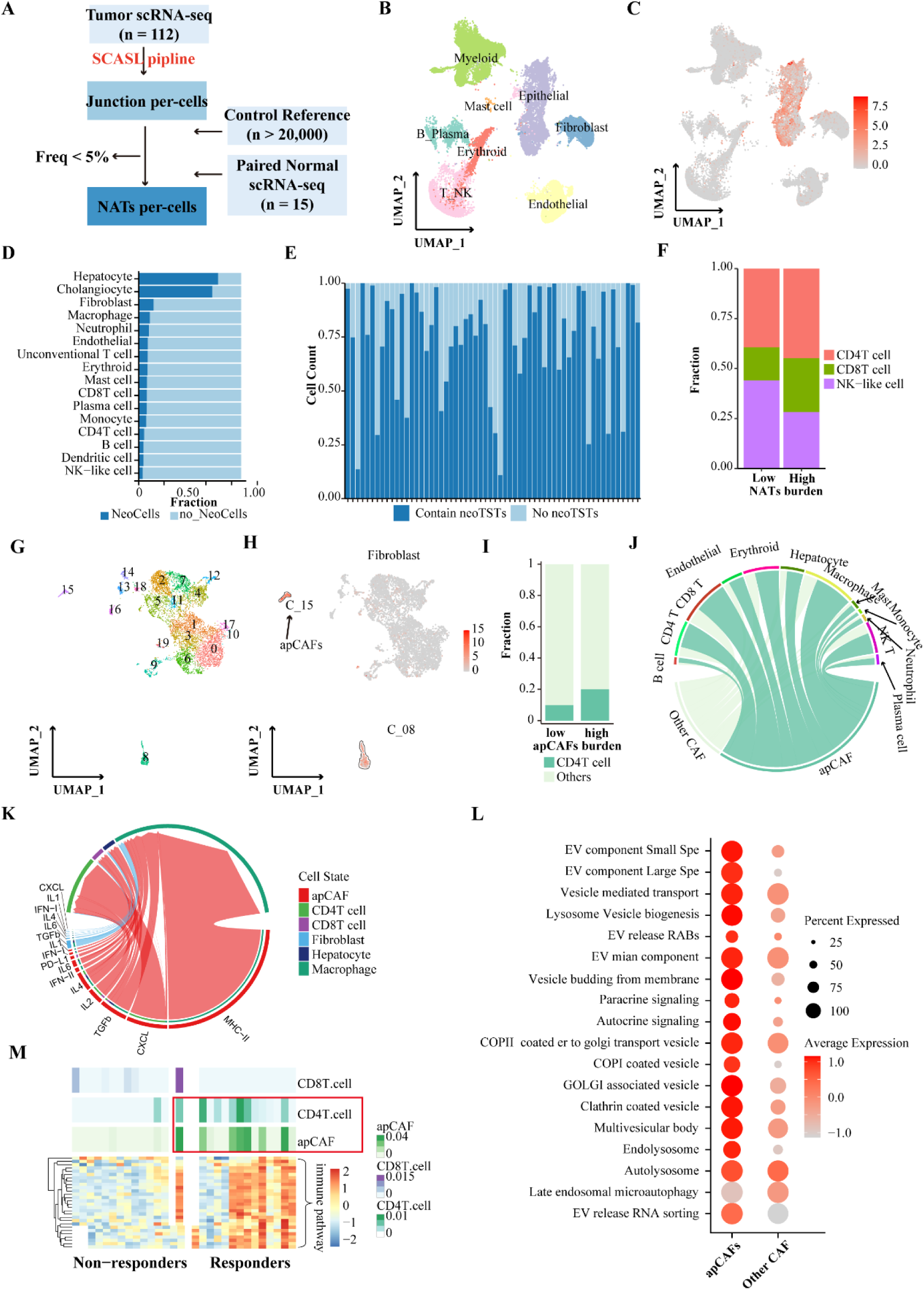
Heterogeneity of neoTSTs at single-cell resolution. (A) Workflow for single-cell level identification of neoTSTs. (B) The UMAP (Uniform Manifold Approximation and Projection) plot displays clustering of single cells (dots) from HRA001748, colored by their cell type annotations. (C) UMAP plot displays the landscape of neoTST count across individual cells in the HCC. (D) Stacked bar plot quantitatively compares the prevalence of neoTSTs among different cell populations in HCC. Each bar represents a distinct cell subtype, with total height normalized to 100%. Dark blue segment: Proportion of cells containing ≥1 neoTST transcript. Light blue segment: neoTST-negative cells. (E) Stacked bar plot quantitatively compares the prevalence of neoTSTs among different samples in HCC. Each bar represents a distinct sample, with total height normalized to 100%. Dark blue segment: Proportion of cells containing ≥1 neoTST. Light blue segment: neoTST-negative cells. (F) Stacked bar plot quantitatively compares the prevalence of T cells between high and low neoTST burden samples. (G) The UMAP plot shows the result of all the re-clustering of cancer associated fibroblasts (CAFs). (H) UMAP plot displays the landscape of neoTST count across individual cells in all CAFs. (I) Stacked bar plot quantitatively compares the prevalence of CD4^+^ T cells between high and low antigen-presenting CAFs (apCAFs) burden samples. (J) Chord diagram, generated through CellChat analysis, maps the signaling crosstalk between apCAFs/other CAFs, and major cell populations in HCC. (K) Chord diagram highlights the immune-modulatory interactions initiated by apCAFs and other CAFs. (L) Dotplot compares the exosome-associated signaling pathway enrichment scores between apCAFs and other CAF subtypes in HCC. (M) Heatmap visualization presents a comprehensive comparison of immune microenvironment features between treatment-responsive (R) and non-responsive (NR) patient groups. Top panel: 4 cellular features (CD4^+^ T cells, CD8^+^ T cells, apCAFs, other CAFs). Bottom panel: 15 key immune pathways (grouped by function). Color gradient: Blue (low), white (medium), red (high) normalized pathway activity.

In addition to hepatocarcinoma cells, neoTSTs were also detected in stromal and immune cells of some patients, with fibroblasts showing the highest abundance (Figure 6D). We extracted cancer-associated fibroblasts (CAFs) from all HCC patients and performed re-clustering analysis. CAFs were grouped into 19 clusters, with neoTSTs predominantly enriched in clusters C_08 and C_15 (Figure 6G, H). Based solely on neoTST expression patterns, fibroblasts were categorized into three groups: C_08, C_15, and others. Cluster C_15 exhibited strong enrichment of neoTSTs, whereas C_08 showed relatively low expression (Supplemental Figure S6A). Previous studies have identified a subset of antigen-presenting CAFs (apCAFs) that express MHC-II molecules and present MHC-II antigens (48). Further annotation revealed that C_15 and C_08 corresponded to apCAFs and myCAFs, respectively (Supplemental Figure S6B). Patient stratification based on apCAF proportion showed that patients with high apCAF levels had significantly increased CD4⁺ T cell infiltration, consistent with MHC-II expression by apCAFs (Figure 6I). Moreover, cell-cell interaction analysis indicated that apCAFs had significantly stronger interactions with CD4⁺ T cells compared to other fibroblasts, primarily involving MHC-II-related pathways (Figure 6J-K, Supplemental Figure S6C). These findings suggest that apCAFs may activate CD4⁺ T cells and promote MHC-II-restricted antigen presentation. Considering that intracellular transcripts may originate from endogenous transcription or exogenous uptake, we analyzed extracellular vesicles (EV)-related pathway activity in both cell types. The majority of EV pathway components showed preferential activation in apCAFs relative to myCAFs. (Figure 6L). These results highlight the potential significance of fibroblasts in immunotherapy.

Given the potential involvement of apCAFs in neoTST generation and presentation, we analyzed sequencing data from 27 HCC patients who received immunotherapy (Responders: 14, Non-responders: 13). Cell-type-specific signatures were constructed based on HCC single-cell RNA sequencing data, and cellular proportions were estimated using CIBERSORT (https://cibersortx.stanford.edu/) (Supplemental Figure S6D). Analysis revealed that the proportion of apCAFs was correlated with the abundance of CD4⁺ T cells in most patients and was significantly enriched in the responder group (Figure 6M). These findings suggest that apCAFs may enhance immunotherapy efficacy by promoting CD4⁺ T cell activity potentially through MHC class II-mediated antigen presentation.

### neoTST elicited antigen-specific CD8^+^ & CD4^+^ T cell responses and inhibited tumor growth in a syngeneic HCC model

To assess the immunogenicity and therapeutic efficacy of neoTST-derived neoantigen vaccines in vivo, we established a murine HCC model using the Hep53.4 cell line (Figure 7A). Through comprehensive genomic profiling, we identified 5 neoTSTs and 20 neoMuts in this system (Figure 7B, this system (Supplemental Figure S7A-B, Supplemental Table S2-4). The neoTST identification pipeline mirrored our human analysis protocol, comprising four key steps: (1) transcriptome assembly, (2) reference database construction, (3) tumor-specific transcript (TST) detection, and (4) neoantigen prediction and validation. For neoMut identification, we detected non-synonymous single nucleotide variants (nsSNVs) and cross-referenced these against murine genomic databases. Candidate neoMuts were subsequently prioritized using MHCPan prediction algorithms, yielding 20 high-confidence targets (Supplemental Table S5). We designed and produced a mixed mRNA-LNP vaccine, which was administered via two intramuscular injections (5 μg/dose) at a 7-day interval to C57BL/6 mice. Flow cytometry analysis demonstrated the superior immunogenicity of neoTSTs, with 60% (3/5, Arl8a, Thebs2, Eps8-neoTSTs) eliciting strong concurrent CD4^+^ and CD8^+^ T-cell responses compared to only 15% (3/20, Spirel, Gtf2i N4bp212-neoMuts) of neoMuts eliciting CD8^+^ T-cell responses (Figure 7B, Supplemental Figure S7C). Notably, the three immunogenic neoTSTs originated from IR events, highlighting non-coding region retention as a critical neoantigen source, and one strongly responsive target (Eps8-neoTST) was driven by a SINE element.

**Figure 7.**
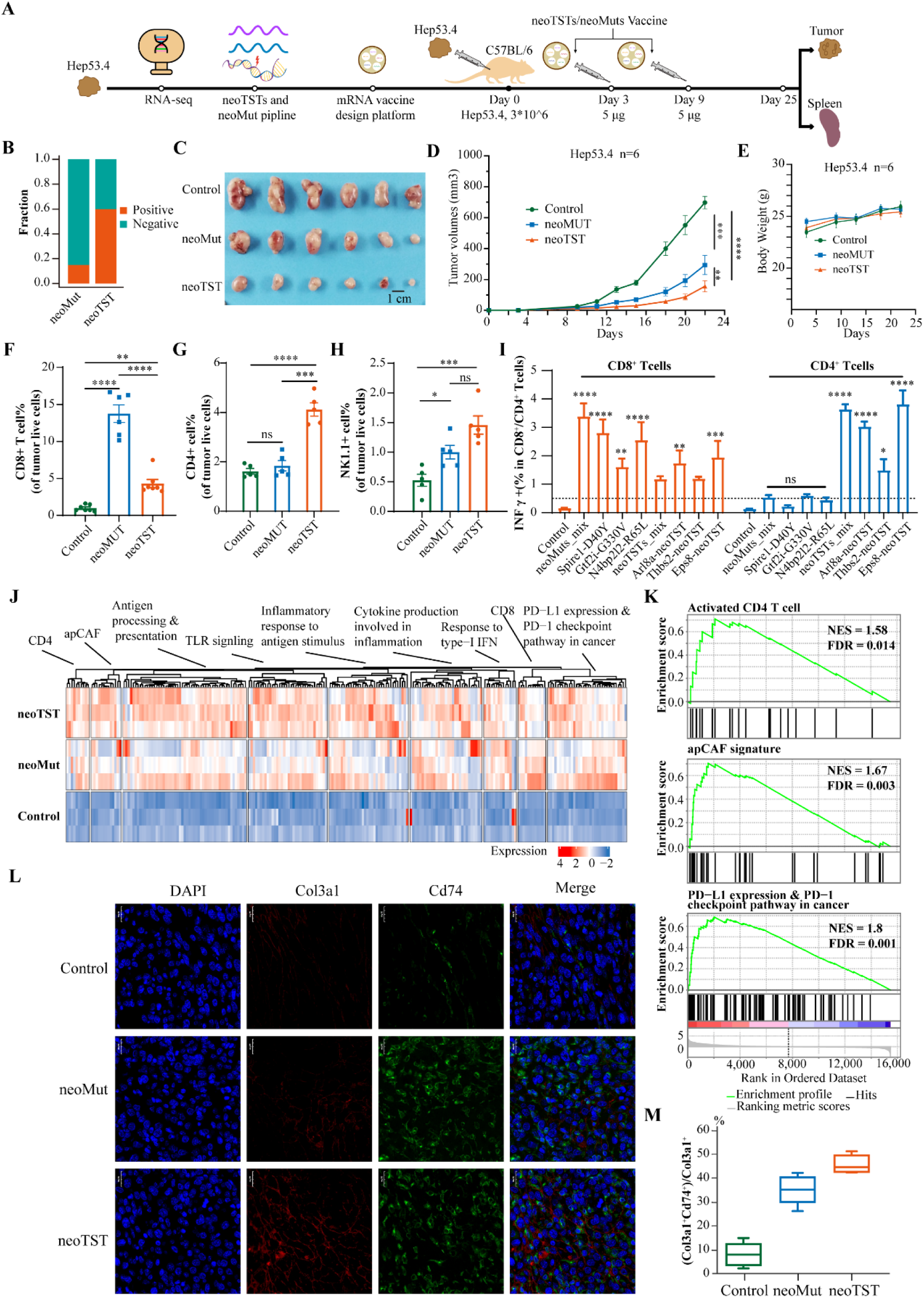
neoTST elicited antigen-specific CD8^+^ & CD4^+^ T cell responses and inhibited tumor growth in a syngeneic HCC model. (A). Schematic of using the murine pancreatic cancer cell line Hep53.4 as a model system to validate in vivo immunogenicity of neoTSTs and assess their therapeutic potential. (B) Stacked bar plot of neoMuts and neoTSTs positivity rates by flow cytometry. (C) Tumor size measurement and visualization. (D) Tumor volume progression (mean ± SEM) over time in different mouse treatment cohorts. (E) Body weight progression (mean ± SEM) over time in different mouse treatment cohorts. (F-H) Cell fraction (mean ± SEM) of CD8^+^ T cell, CD4^+^ T cell and NK T cell in different mouse treatment cohorts. (I) Proportions of CD8^+^ T cells (left) and CD4^+^ T cells (right) induced by neoTST versus neoMut neoantigens. One-way ANOVA was used; data shown as mean± SEM. *P<0.05, **P<0.01, ***P<0.001. Control: unvaccinated mice. (J) Clustered heatmap compares the immune microenvironment profiles of different treatment groups in the mouse model, integrating immunity pathway and cell-type-specific genes expression. (K) Gene Set Enrichment Analysis (GSEA) results comparing neoTST-treated versus control mice, focusing on three key biological modules: Activated CD4^+^ T Cell (top, NES=1.58, FDR=0.014); apCAF Signature (middle, NES=1.67, FDR=0.003); PD-L1 expression and PD-l checkpoint pathway in cancer (bottom, NES=1.8, FDR=0.001). (L) Representative IF staining images of Col3a1 (red) and Cd74 (green) in the tumors from indicated groups. (M) Fraction of apCAF in CAFs ((Col3a1^+^Cd74^+^)/Col3a1^+^).

Furthermore, mice bearing subcutaneous Hep53.4 tumors were vaccinated with the respective neoantigen vaccines on days 5 and 12 post-tumor inoculation. Tumor growth was monitored, and after 21 days, subcutaneous tumors and spleens were harvested. Results showed that both mutation-derived and TST-derived neoantigens effectively suppressed tumor growth, with neoTSTs demonstrating superior efficacy (Figure 7C-D). Body weight monitoring indicated no significant systemic toxicity, suggesting favorable safety profiles of the vaccines (Figure 7G). Comparative analysis of tumor-infiltrating lymphocytes revealed distinct immune activation patterns between vaccination groups. Mice immunized with neoMuts exhibited significantly higher CD8^+^ T cell infiltration compared to both control (p <0.001) and neoTSTs (p <0.01) groups (Figure 7F). Notably, while neoMuts vaccination showed no significant effect on CD4^+^ T cell recruitment, neoTSTs immunization induced robust CD4^+^ T cell infiltration (p <0.001 versus control), demonstrating their unique capacity to activate both MHC class I and II antigen presentation pathways (Figure 7G). Both vaccine groups showed increased NK cell infiltration relative to controls (p <0.05), though the difference between neoMuts and neoTSTs groups did not reach statistical significance (Figure 7H). These results indicate that neoTSTs can simultaneously activate the immune pathways of both MHC I and MHC II, and they exhibit a stronger cytotoxic potential compared to neoMuts. Following independent transfection of the three neoMuts and three neoTSTs, flow cytometric analysis revealed distinct immunogenic profiles. While both vaccine types elicited antigen-specific CD8^+^ T cell responses, only neoTSTs demonstrated the capacity to induce robust CD4^+^ T cell activation (Figure 7I). These findings provide direct experimental evidence that neoTSTs uniquely engage both MHC class I and II antigen presentation pathways, establishing their superior therapeutic potential compared to neoMuts for comprehensive anti-tumor immunity.

### neoTST enhances the infiltration of apCAF in a syngeneic HCC model

To further compare the effects of neoantigen vaccines, we collected Hep53.4 tumor tissues from mice that were either unvaccinated or vaccinated with neoMuts- or neoTSTs-based vaccines for RNA sequencing. Transcriptomic profiling revealed that both neoantigen vaccines robustly activated hallmark immune pathways, including antigen processing and presentation, cytokine production involved in inflammation, PD-L1 expression and PD-1 checkpoint pathway in cancer, inflammatory response to antigen stimulus, response to type-I IFN, TLR signaling (Figure 7J). Additionally, both vaccines potently upregulated gene signatures associated with Activated CD4^+^ T cells, Activated CD8^+^ T cells, and apCAF, with the neoTST vaccine demonstrating stronger activation than the neoMut vaccine. GSEA analysis further revealed distinct response patterns: while neoTST-vaccinated mice showed significant enrichment of Activated CD4^+^ T cell (NES = 1.58, FDR = 0.014), apCAF signatures (NES = 1.67, FDR = 0.003), and PD-L1 expression and PD-1 checkpoint pathway compared to controls (NES = 1.8, FDR = 0.001), only PD-L1 expression and PD-1 checkpoint pathway was markedly activated in neoMut-treated mice (NES = 1.70, FDR = 0.01) (Figure 7K, Supplemental Table S6-7). This also supported that neoTSTs may promote the CD4^+^ T cell immune pathway through apCAFs.

Quantitative immunofluorescence revealed significantly enhanced apCAF (Col3a1^+^Cd74^+^) infiltration in neoTST-vaccinated tumors compared to control, whereas neoMut-vaccinated tumors showed intermediate infiltration level (Figure 7L-M), confirming the unique capacity of neoTST vaccines to remodel the tumor stroma through apCAF mobilization. These findings demonstrate that neoTSTs may exhibit superior antigen-presenting capacity compared to neoMuts. Furthermore, neoantigen vaccine stimulation robustly activates diverse immune pathways, driving the conversion of immunologically “cold” tumors into “hot” ones. This transformation creates a permissive microenvironment for enhanced efficacy when combined with ICIs, offering a promising combinatorial immunotherapy strategy.

## DISCUSSION

Our study systematically identified an average of 60 neoTSTs per patient across 1,013 liver cancers through a stringent analytical pipeline. These neoTSTs predominantly originated from aberrant splicing events and intron retention, driven by TEs and the transcription factor HNF4A. Interestingly, besides tumor cells, apCAFs were also found to produce a subset of neoTSTs, which primarily encode MHC-II-restricted epitopes and facilitate CD4⁺ T cell activation. Functional validation in an HLA-A transgenic model confirmed the ability of neoTSTs to elicit specific T cell responses, and further experiments in HCC mouse models demonstrated their superior capacity in activating antitumor immunity and suppressing tumor growth. Collectively, our work identifies a large repertoire of therapeutically promising neoTSTs in HCC, offering novel targets and insights to advance immunotherapy and therapeutic development for liver cancer.

In recent years, growing attention has been drawn to the importance of transcript-derived neoantigens in cancer patients. Previous studies have shown that analyzing alternative splicing enables more accurate identification of transcript isoforms (27). Conventional approaches for assessing transcript splicing, such as Percent Spliced In (PSI) analysis, primarily quantify splicing events but are unable to directly identify novel splicing junctions and rely heavily on existing annotations. Recently, several tools-such as SPLICE-neo (49), SNAF (50), and IRIS (51) -have been developed to screen immunotherapeutic targets by detecting specific splicing sites. However, these tools predominantly focus on spliced events and often overlook transcripts without splicing. Moreover, many studies rely solely on splicing references from resources like GTEx Portal, which may limit the ability to comprehensively capture tumor-specific transcripts and introduce false positives. Additionally, we introduce a novel workflow that integrates RNA-seq data from diverse tissues, populations, and platforms to construct a comprehensive reference dataset. Through multi-level comparisons, our approach identifies both multi-exon and single-exon neoTSTs, thereby expanding the repertoire of detectable neoantigens and improving prediction accuracy. Previous research suggests that shorter open reading frames (ORFs) facilitate neoantigen processing and presentation, indicating that single-exon neoTSTs may possess stronger immunogenic potential (40, 41). We also designed a unique alignment strategy: a hash-formatted reference peptide pool was built from known protein sequences, followed by pre-alignment of full-length neoTST protein sequences to identify novel specific peptides before further neoantigen prediction. This method reduces false positives while retaining a broad spectrum of potential neoantigens and significantly accelerates computational processing. Furthermore, unlike conventional methods that apply differential expression thresholds (e.g., IRIS) or global fold-change filters (e.g., SNAF), we established a comprehensive reference dataset encompassing three biological categories: 30 normal tissue types, matched adjacent non-tumor tissues, and non-cancer disease specimens. Each layer underwent independent benchmarking before feature integration. This multi-tiered comparison enables precise identification of tumor-specific splicing events, avoiding signal dilution caused by mixed tissue types or physiological states, thereby substantially enhancing neoantigen specificity. Notably, during the finalization of our study, a contemporaneous study identified transcript-derived neoantigens in HCC through alternative splicing analysis (52); however, this work was limited to comparisons with only adjacent normal liver tissues, exhibited a pronounced false-positive rate, and crucially, did not address the epigenetic-driven transcriptional variants identified in our work. Overall, our novel workflow improves the accuracy of transcript-derived neoantigen identification and expands the candidate target pool for liver cancer.

In contrast to the highly individualized neoantigens derived from SNVs, neoTSTs originate from distinct molecular mechanisms-namely, aberrant splicing events and chromatin accessibility alterations. This unique origin confers two key advantages to neoTSTs. First, their splicing patterns exhibit reproducibility, enabling the possibility of shared neoTSTs across different liver cancer patients (i.e., “public neoantigens”). This characteristic opens avenues for developing universal T-cell therapies or vaccines, thereby overcoming the limitations of personalized mutation-based neoantigen approaches. Second, previous studies have revealed extensive de novo chromatin opening events in tumors, with these newly accessible regions often enriched for transcription factor binding sites (53–55). Our finding further demonstrates that HCC exhibits widespread specific open chromatin regions that are not only enriched with binding sites for transcription factors such as HNF4A-which shows a significant positive correlation with neoTST burden-but also activate cryptic regulatory elements (e.g., transposable elements), leading to the production of numerous F-neoTSTs utilizing novel transcription start sites. From a structural and functional perspective, neoTSTs exhibit multiple therapeutic implications: 1) Membrane localization: Approximately 15% of neoTSTs contain conserved or de novo transmembrane domains, facilitating efficient presentation via MHC class I/II molecules and enhancing T-cell recognition. This surface-exposed feature also makes them ideal targets for antibody-based therapies, such as bispecific antibodies or CAR-T. 2) Dual mechanisms of action: neoTSTs may directly modulate tumor biology through retained functional domains (e.g., kinase domains), as illustrated in Figure 5SC. These mechanisms include sustained activation of pro-proliferative signaling pathways (when kinase domains are preserved) or loss of tumor-suppressive functions (due to altered subcellular localization).

It is noteworthy that while current neoantigen research primarily focuses on antigens presented by HLA class I molecules and their role in activating CD8⁺ T cells, accumulating evidence highlights the critical function of HLA class II molecules and the CD4⁺ T cell responses they mediate in enhancing antitumor immunity (56). Through in-depth analysis of single-cell sequencing and RNA-seq data from immunotherapy patients, we observed a significant correlation between the activation status of CD4⁺ T cells and therapeutic efficacy, a process potentially under fine-tuned regulation by apCAFs. Animal studies further confirmed that neoTSTs not only effectively activate CD8⁺ T cells but also concurrently stimulate CD4⁺ T cells via the HLA class II pathway, thereby exerting synergistic antitumor effects. Although the accuracy of current HLA class II antigen prediction tools remains to be improved, our study employed a non-restrictive HLA-I screening strategy, which retains a broader spectrum of potential epitopes and effectively encompasses candidate targets likely presented through the HLA-II pathway, thus providing methodological support for comprehensively exploring the immunotherapeutic potential of neoTSTs. Animal experiments revealed that 60% of neoTSTs could simultaneously stimulate both CD4⁺ and CD8⁺ T cell responses. However, NetMHCpan predictions indicated that some of these neoTSTs lack predicted HLA-II binding motifs, further underscoring the necessity of a non-restrictive HLA screening approach. These findings not only expand our understanding of tumor immune response mechanisms but also provide a critical theoretical foundation for developing novel combination immunotherapy strategies that concurrently target both CD8⁺ and CD4⁺ T cells.

While this study establishes a comprehensive HCC neoantigen repository and validates selected neoTSTs in preclinical models, several limitations should be acknowledged. First, although we confirmed the immunogenicity and antitumor activity of representative neoTSTs, the majority of candidates remain experimentally unverified, necessitating systematic functional characterization to assess their potential as shared therapeutic targets. Second, our immunopeptidomic validation, while informative, was constrained by the limited sample size and lack of paired tumor-normal data from existing datasets, highlighting the need for larger-scale, matched analyses. Most importantly, current immunogenicity prediction tools exhibit marked bias toward MHC-I epitopes due to historical data limitations, while our findings demonstrate neoTSTs’ pronounced MHC-II immunogenicity-a critical but understudied aspect of antitumor immunity. These results underscore the urgent need for: (1) MHC-II-specific experimental validation platforms, and (2) development of balanced prediction algorithms incorporating both MHC-I and -II immunogenicity data to fully exploit neoTSTs’ therapeutic potential.

## CONCLUSION

Our study established a comprehensive repository of potential neoTST candidates in liver cancer and elucidated their distinct splicing features and underlying drivers. These findings expand the understanding of neoantigen sources in liver cancer, providing a robust pool of high-potential targets that may enable shared therapeutic strategies. This resource offers critical support for enhancing liver cancer immunotherapy efficacy. Experimental validation-both *in vitro* and *in vivo*--confirmed the substantial immunotherapeutic potential of selected neoTSTs and their advantages in improving therapeutic outcomes. Collectively, this work holds considerable promise for broadening the clinical applicability of neoantigen-based immunotherapies.

## DATA AVAILABILITY STATEMENT

All RNA-seq data of tumor are available at the Gene Expression Omnibus (GSE112221, GSE144269, GSE94660, GSE114564, GSE124535, GSE140462, GSE148355, GSE77314, GSE77509, GSE104766, GSE133039, GSE151347, GSE81928, GSE89775, GSE107943, GSE119336, GSE162396, GSE63420). Proteomic raw data (PDC000198) was retrieved from the Proteomic Data Commons (PDC). Immunopeptidome raw datra (PXD023143) was download from Proteome Xchange (proteomecentral.proteomexchange.org). TCGA clinical metadata was acquired from Thorsson etal. with HLA typing data from the Pan-CancerAtlas (https://gdc.cancer.gov/aboutdata/publications/panimmune). Somatic mutationannotations (MAF files) were downloaded from the TCGA-LIHC repository(https://portal.gdc.cancer.gov/); somatic mutation-derived neoantigens (SNVs) forTCGA were obtained from TSNAdb (https://pgx.zju.edu.cn/tsnadb/download/). Single-cell RNA-seq data (HRA001748) was apply from GSA-Human (https://ngdc.cncb.ac.cn/gsa-human. RepeatMasker annotations (hg38) were sourced from the UCSC Genome Browser. Other data are available on reasonable request.

## AUTHOR CONTRIBUTIONS

Shenglin Huang and Linhui Liang conceptualized the study and designed the experiments. Peng Lin conducted the data analysis and interpretation. Yifan Wen conducted animal experiments and acquisition of data. Feifei Zhang and Yaoming Su conducted cell experiments. Jingjing Zhao, Zhixiang Hu and Yan Li contributed to the interpretation of the results. Peng Lin and Shenglin Huang wrote the original paper. Jingjing Zhao, Hongwu Yu, Zhuting Fang helped in editing the paper. All authors read and approved the final manuscript.

## ACKNOWLEDGMENTS

The authors thank the patients and investigators who participated in the publicly available data sets for providing data.

## CONFLICTS OF INTEREST

All the authors have no reported conflicts of interest.

**Footnotes**

## Abbreviations

TST: Tumor-specific transcript
neoTST: neoantigen-encoding tumor-specific transcripts
neoMut: mutation-derived neoantigen
TMD: transmembrane domain
ORF: Open read frame
TE: Transposable elements
TSS: Transcriptional start site
EV: Extracellular vesicle
ApCAF: Antigen-presenting cancer-associated fibroblast

## Supplemental Figure

**Supplemental Figure S1.** Landscape of Neoantigen-encoding tumor-specific transcripts (neoTSTs) in 1,013 Liver Cancer. (A) Bar plot provides a comprehensive breakdown of the biological specimens analyzed in the study, categorized by tissue origin and disease status. (B) Quantifies splice junction of controls from raw BAM files (left, n > 3,000) and junction frequency threshold: ≥5% occurrence in any GTEX tissue (right, n > 20,000). (C) Total number of RNA splicing junctions detected per tumor sample, stratified by molecular subtypes of liver cancer. (D) Prevalence of multi-exonic transcripts (METs) and single-exonic transcripts (SETs) count in different cancer types. X-axis: Cancer types (HB, HCC, ICC). Y-axis: neoTSTs count. Orange dashed: Mean SETs frequency (16 events/sample). Green dashed: Mean MET alteration frequency (44 events/sample). (E) Venn diagram quantifies neoTSTs identified in 3 major liver cancer types, revealing both shared and specific neoTSTs count. (F) Prevalence of hepatitis B virus (HBV) infection in the study’s HCC samples. (G) Count of neoTSTs between HBV-positive and HBV-negative HCC patients. Statistical significance was determined by Wilcoxon rank-sum test (***p < 0.001). (H) Kaplan-Meier survival analysis stratified by neoTST burden (high vs. low) of different HLA subtype in TCGA-LIHC cohort. Log-rank test was used. The optimal cutoff point of TST-derived neoantigen burden was applied. (I) Kaplan-Meier survival analysis stratified by neoTST burden (high vs. low) of different HLA subtype in in-house cohort. Log-rank test was used. The optimal cutoff point of TST-derived neoantigen burden was applied.

**Supplemental Figure S2.** Mass Spectrometry and Immunopeptidomics-Based Identification and Immunogenicity Assessment of neoTSTs in Liver Cancer. (A) Distribution pattern of the 20 neoMuts across HCC samples from TCGA-LIHC. (B) Proteomic identification of neoTSTs versus canonical proteins in HCC. (C) Immunopeptidomic identification of neoTSTs in HCC. (D) The number of different HLA subtypes identified by the neoTSTs in the immunopeptide spectrum. The HLA subtypes with the highest frequencies are highlighted.

**Supplemental Figure S3.** Retention of noncoding sequences and transposon-driven activation generate high-yield neoantigen expansion. (A-B) Classification and proportions of splicing junction types. (C) Distribution of splicing signals from different sources. (D-E) Proportions of Chimeric-neoTSTs and nonchimeric-neoTSTs driven by J or neoJ. (F) Examines the spatial distribution of predicted neoantigen-peptides (neoPeps) across tumor-specific transcript isoforms (neoTSTs) based on their protein sequence positions. Here, 0 indicates that neoPep is located at the beginning of the neoTST protein. 1-10 indicate that neoPep is located at the n/10 region of the neoTST protein. (G) The boxplot compares the expression levels of individual TE subfamilies between high neoTSTs burden samples and low neoTSTs burden samples. Each box represents the distribution of subfamily expression within a TE type. Dots indicate outliers. (H) Scatter plot showing the positive correlation between log2(LTR7Y expression) (x-axis) and neoTST burden (y-axis, count per sample). A trendline (blue dashed line) indicates a significant linear relationship (r = 0.27, p < 0.001). (I) The quantity of neoPeps derived from TE. (J) The frequencies of the top 20 TE-neoTST distributions and the frequencies, quantities, and MHC affinity scores of the resulting neoPeps.

**Supplemental Figure S4.** neoTSTs could generate de novo transmembrane domains (TMDs). (A) The figure presents the 3D structural models of: Wild-type ZPLD1 (blue ribbon diagram) ZPLD1-neoTST (red ribbon diagram); Wild-type GJA10 (blue ribbon diagram) GJA10-neoTST (red ribbon diagram); Wild-type HERC4 (blue ribbon diagram) HERC4-neoTST (red ribbon diagram).

**Supplemental Figure S5.** HNF4A binding activates neoTSTs transcription in HCC. (A) The UMAP plot highlights HNF4A expression within the single-cell landscape of liver cancer.

**Supplemental Figure S6.** Heterogeneity of neoTSTs at single-cell resolution. (A) Count of neoTST in C_15, C_08 and other cancer-associated fibroblasts (CAFs). (B) Signatures expression of antigen-presenting CAF (apCAF) in C_15, C_08 and other CAFs. (C) KEGG enrichment results of up regulated genes in C_15, only highlights and marks the antigen processing and presentation pathway. (D) Heatmap shows the expression profiles of canonical marker genes across distinct cell populations.

**Supplemental Figure S7.** neoTST elicited antigen-specific CD8+ & CD4+ T cell responses and inhibited tumor growth in a syngeneic HCC model. (A) Characterization of representative neoTSTs in Hep53.4. (B) Sashimi plot of 3 neoTSTs in Hep53.4: Arl8a-neoTST, Thbs2-neoTST and Eps8-neoTST. (C) Flow cytometry results of 20 neoMut and 5 neoTST vaccines.

## REFERENCES

1. Bray F, Laversanne M, Sung H, Ferlay J, Siegel RL, Soerjomataram I, Jemal A. Global cancer statistics 2022: GLOBOCAN estimates of incidence and mortality worldwide for 36 cancers in 185 countries. CA Cancer J Clin 2024;74:229–263.

2. Forner A, Reig M, Bruix J. Hepatocellular carcinoma. Lancet 2018;391:1301–1314.

3. Du JS, Hsu SH, Wang SN. The Current and Prospective Adjuvant Therapies for Hepatocellular Carcinoma. Cancers (Basel) 2024;16.

4. Kolamunnage-Dona R, Berhane S, Potts H, Williams EH, Tanner J, Janowitz T, Hoare M, et al. Sorafenib is associated with a reduced rate of tumour growth and liver function deterioration in HCV-induced hepatocellular carcinoma. J Hepatol 2021;75:879–887.

5. Kudo M, Ren Z, Guo Y, Han G, Lin H, Zheng J, Ogasawara S, et al. Transarterial chemoembolisation combined with lenvatinib plus pembrolizumab versus dual placebo for unresectable, non-metastatic hepatocellular carcinoma (LEAP-012): a multicentre, randomised, double-blind, phase 3 study. Lancet 2025;405:203–215.

6. Zhou J, Bai L, Luo J, Bai Y, Pan Y, Yang X, Gao Y, et al. Anlotinib plus penpulimab versus sorafenib in the first-line treatment of unresectable hepatocellular carcinoma (APOLLO): a randomised, controlled, phase 3 trial. Lancet Oncol 2025;26:719–731.

7. Shen KY, Zhu Y, Xie SZ, Qin LX. Immunosuppressive tumor microenvironment and immunotherapy of hepatocellular carcinoma: current status and prospectives. J Hematol Oncol 2024;17:25.

8. Li B, Li Y, Zhou H, Xu Y, Cao Y, Cheng C, Peng J, et al. Multiomics identifies metabolic subtypes based on fatty acid degradation allocating personalized treatment in hepatocellular carcinoma. Hepatology 2024;79:289–306.

9. Ren Z, Xu J, Bai Y, Xu A, Cang S, Du C, Li Q, et al. Sintilimab plus a bevacizumab biosimilar (IBI305) versus sorafenib in unresectable hepatocellular carcinoma (ORIENT-32): a randomised, open-label, phase 2-3 study. Lancet Oncol 2021;22:977–990.

10. Finn RS, Qin S, Ikeda M, Galle PR, Ducreux M, Kim TY, Kudo M, et al. Atezolizumab plus Bevacizumab in Unresectable Hepatocellular Carcinoma. N Engl J Med 2020;382:1894–1905.

11. De Martin E, Fulgenzi CAM, Celsa C, Laurent-Bellue A, Torkpour A, Lombardi P, D’Alessio A, et al. Immune checkpoint inhibitors and the liver: balancing therapeutic benefit and adverse events. Gut 2025;74:1165–1177.

12. Yang Y, Ni Q, Li H, Sun J, Zhou X, Qu L, Wang L, et al. Genomic and the tumor microenvironment heterogeneity in multifocal hepatocellular carcinoma. Hepatology 2025;82:582–598.

13. Benoit A, Vogin G, Duhem C, Berchem G, Janji B. Lighting Up the Fire in the Microenvironment of Cold Tumors: A Major Challenge to Improve Cancer Immunotherapy. Cells 2023;12.

14. Chen P, Chen D, Bu D, Gao J, Qin W, Deng K, Ren L, et al. Dominant neoantigen verification in hepatocellular carcinoma by a single-plasmid system coexpressing patient HLA and antigen. J Immunother Cancer 2023;11.

15. Blass E, Ott PA. Advances in the development of personalized neoantigen-based therapeutic cancer vaccines. Nat Rev Clin Oncol 2021;18:215–229.

16. Chen H, Li Z, Qiu L, Dong X, Chen G, Shi Y, Cai L, et al. Personalized neoantigen vaccine combined with PD-1 blockade increases CD8(+) tissue-resident memory T-cell infiltration in preclinical hepatocellular carcinoma models. J Immunother Cancer 2022;10.

17. Lang F, Schrörs B, Löwer M, Türeci Ö, Sahin U. Identification of neoantigens for individualized therapeutic cancer vaccines. Nat Rev Drug Discov 2022;21:261–282.

18. Peng S, Chen S, Hu W, Mei J, Zeng X, Su T, Wang W, et al. Combination Neoantigen-Based Dendritic Cell Vaccination and Adoptive T-Cell Transfer Induces Antitumor Responses Against Recurrence of Hepatocellular Carcinoma. Cancer Immunol Res 2022;10:728–744.

19. Xie N, Shen G, Gao W, Huang Z, Huang C, Fu L. Neoantigens: promising targets for cancer therapy. Signal Transduct Target Ther 2023;8:9.

20. Dolina JS, Lee J, Brightman SE, McArdle S, Hall SM, Thota RR, Zavala KS, et al. Linked CD4+/CD8+ T cell neoantigen vaccination overcomes immune checkpoint blockade resistance and enables tumor regression. J Clin Invest 2023;133.

21. Weber JS, Carlino MS, Khattak A, Meniawy T, Ansstas G, Taylor MH, Kim KB, et al. Individualised neoantigen therapy mRNA-4157 (V940) plus pembrolizumab versus pembrolizumab monotherapy in resected melanoma (KEYNOTE-942): a randomised, phase 2b study. Lancet 2024;403:632–644.

22. Berraondo P, Cuesta R, Sanmamed MF, Melero I. Immunogenicity and Efficacy of Personalized Adjuvant mRNA Cancer Vaccines. Cancer Discov 2024;14:2021–2024.

23. Yarchoan M, Gane EJ, Marron TU, Perales-Linares R, Yan J, Cooch N, Shu DH, et al. Personalized neoantigen vaccine and pembrolizumab in advanced hepatocellular carcinoma: a phase 1/2 trial. Nat Med 2024;30:1044–1053.

24. Rojas LA, Sethna Z, Soares KC, Olcese C, Pang N, Patterson E, Lihm J, et al. Personalized RNA neoantigen vaccines stimulate T cells in pancreatic cancer. Nature 2023;618:144–150.

25. Sethna Z, Guasp P, Reiche C, Milighetti M, Ceglia N, Patterson E, Lihm J, et al. RNA neoantigen vaccines prime long-lived CD8(+) T cells in pancreatic cancer. Nature 2025;639:1042–1051.

26. Hu Z, Guo X, Li Z, Meng Z, Huang S. The neoantigens derived from transposable elements - A hidden treasure for cancer immunotherapy. Biochim Biophys Acta Rev Cancer 2024;1879:189126.

27. Li Q, Li Z, Chen B, Zhao J, Yu H, Hu J, Lai H, et al. RNA splicing junction landscape reveals abundant tumor-specific transcripts in human cancer. Cell Rep 2024;43:114893.

28. Jayasinghe RG, Cao S, Gao Q, Wendl MC, Vo NS, Reynolds SM, Zhao Y, et al. Systematic Analysis of Splice-Site-Creating Mutations in Cancer. Cell Rep 2018;23:270–281.e273.

29. Sun Y, Xiong B, Shuai X, Li J, Wang C, Guo J, Cheng Z, et al. Downregulation of HNRNPA1 induced neoantigen generation via regulating alternative splicing. Mol Med 2024;30:85.

30. Frankiw L, Baltimore D, Li G. Alternative mRNA splicing in cancer immunotherapy. Nat Rev Immunol 2019;19:675–687.

31. Zhao J, Li Q, Li Y, He X, Zheng Q, Huang S. ASJA: A Program for Assembling Splice Junctions Analysis. Comput Struct Biotechnol J 2019;17:1143–1150.

32. Li Q, Lai H, Li Y, Chen B, Chen S, Li Y, Huang Z, et al. RJunBase: a database of RNA splice junctions in human normal and cancerous tissues. Nucleic Acids Res 2021;49:D201–d211.

33. Guo W, Hu Z, Bao Y, Li Y, Li S, Zheng Q, Lyu D, et al. A LIN28B Tumor-Specific Transcript in Cancer. Cell Rep 2018;22:2016–2025.

34. Yang YS, Jin X, Li Q, Chen YY, Chen F, Zhang H, Su Y, et al. Superenhancer drives a tumor-specific splicing variant of MARCO to promote triple-negative breast cancer progression. Proc Natl Acad Sci U S A 2022;119:e2207201119.

35. Dobin A, Davis CA, Schlesinger F, Drenkow J, Zaleski C, Jha S, Batut P, et al. STAR: ultrafast universal RNA-seq aligner. Bioinformatics 2013;29:15–21.

36. Pertea M, Pertea GM, Antonescu CM, Chang TC, Mendell JT, Salzberg SL. StringTie enables improved reconstruction of a transcriptome from RNA-seq reads. Nat Biotechnol 2015;33:290–295.

37. Wang L, Park HJ, Dasari S, Wang S, Kocher JP, Li W. CPAT: Coding-Potential Assessment Tool using an alignment-free logistic regression model. Nucleic Acids Res 2013;41:e74.

38. Orenbuch R, Filip I, Comito D, Shaman J, Pe’er I, Rabadan R. arcasHLA: high-resolution HLA typing from RNAseq. Bioinformatics 2020;36:33–40.

39. Reynisson B, Alvarez B, Paul S, Peters B, Nielsen M. NetMHCpan-4.1 and NetMHCIIpan-4.0: improved predictions of MHC antigen presentation by concurrent motif deconvolution and integration of MS MHC eluted ligand data. Nucleic Acids Res 2020;48:W449–w454.

40. Thompson MK, Rojas-Duran MF, Gangaramani P, Gilbert WV. The ribosomal protein Asc1/RACK1 is required for efficient translation of short mRNAs. Elife 2016;5.

41. Wu Q, Wright M, Gogol MM, Bradford WD, Zhang N, Bazzini AA. Translation of small downstream ORFs enhances translation of canonical main open reading frames. Embo j 2020;39:e104763.

42. Demircioğlu D, Cukuroglu E, Kindermans M, Nandi T, Calabrese C, Fonseca NA, Kahles A, et al. A Pan-cancer Transcriptome Analysis Reveals Pervasive Regulation through Alternative Promoters. Cell 2019;178:1465–1477.e1417.

43. Karttunen K, Patel D, Xia J, Fei L, Palin K, Aaltonen L, Sahu B. Transposable elements as tissue-specific enhancers in cancers of endodermal lineage. Nat Commun 2023;14:5313.

44. Jang HS, Shah NM, Du AY, Dailey ZZ, Pehrsson EC, Godoy PM, Zhang D, et al. Transposable elements drive widespread expression of oncogenes in human cancers. Nat Genet 2019;51:611–617.

45. Chuong EB, Elde NC, Feschotte C. Regulatory evolution of innate immunity through co-option of endogenous retroviruses. Science 2016;351:1083–1087.

46. Chuong EB, Rumi MA, Soares MJ, Baker JC. Endogenous retroviruses function as species-specific enhancer elements in the placenta. Nat Genet 2013;45:325–329.

47. Li Z, Zhang H, Li Q, Feng W, Jia X, Zhou R, Huang Y, et al. GepLiver: an integrative liver expression atlas spanning developmental stages and liver disease phases. Sci Data 2023;10:376.

48. Elyada E, Bolisetty M, Laise P, Flynn WF, Courtois ET, Burkhart RA, Teinor JA, et al. Cross-Species Single-Cell Analysis of Pancreatic Ductal Adenocarcinoma Reveals Antigen-Presenting Cancer-Associated Fibroblasts. Cancer Discov 2019;9:1102–1123.

49. Wickland DP, McNinch C, Jessen E, Necela B, Shreeder B, Lin Y, Knutson KL, et al. Comprehensive profiling of cancer neoantigens from aberrant RNA splicing. J Immunother Cancer 2024;12.

50. Li G, Mahajan S, Ma S, Jeffery ED, Zhang X, Bhattacharjee A, Venkatasubramanian M, et al. Splicing neoantigen discovery with SNAF reveals shared targets for cancer immunotherapy. Sci Transl Med 2024;16:eade2886.

51. Pan Y, Phillips JW, Zhang BD, Noguchi M, Kutschera E, McLaughlin J, Nesterenko PA, et al. IRIS: Discovery of cancer immunotherapy targets arising from pre-mRNA alternative splicing. Proc Natl Acad Sci U S A 2023;120:e2221116120.

52. Zhao H, Cheng Y, Zhang T, Wang Q, Xu Y, Wang G, Song Y, et al. Harnessing alternative splicing for off-the-shelf mRNA neoantigen vaccines in hepatocellular carcinoma. Cell Res 2025.

53. Corces MR, Granja JM, Shams S, Louie BH, Seoane JA, Zhou W, Silva TC, et al. The chromatin accessibility landscape of primary human cancers. Science 2018;362.

54. Britton E, Rogerson C, Mehta S, Li Y, Li X, Fitzgerald RC, Ang YS, et al. Open chromatin profiling identifies AP1 as a transcriptional regulator in oesophageal adenocarcinoma. PLoS Genet 2017;13:e1006879.

55. Xia L, Lu J, Qin Y, Huang R, Kong F, Deng Y. Analysis of chromatin accessibility in peripheral blood mononuclear cells from patients with early-stage breast cancer. Front Pharmacol 2024;15:1465586.

56. Brightman SE, Naradikian MS, Miller AM, Schoenberger SP. Harnessing neoantigen specific CD4 T cells for cancer immunotherapy. J Leukoc Biol 2020;107:625–633.

